# Anomaly detection uncovers emerging novelty in coral assemblages of the Great Barrier Reef

**DOI:** 10.64898/2026.10.15.759496

**Authors:** Timothy L Staples, Sun Wook Kim, Kerryn Crossman, John M Pandolfi

## Abstract

Earth’s biosphere is being altered by human actions, with all ecosystems, especially coral reefs, rapidly transforming in response to climate change and local stressors. As these changes manifest, ecology needs tools to measure change from past states, termed “ecological novelty”, within complex multidimensional data such as taxa-abundance time series. Here we use an anomaly detection method new to ecology, Local Outlier Factor (LOF), to measure ecological novelty via neighborhood density in compositional space. We estimated novelty using LOF within 106 coral time series, spanning 29 years, from a long-term monitoring program on the Great Barrier Reef (GBR). We tested whether reef assemblages were becoming more novel over time and whether novelty correlated with common ecological indicators: coral cover, diversity and assemblage turnover. We found novelty correlated with low coral cover, low diversity and high turnover. Even after accounting for these correlations, we found reefs became more novel over time, with novelty across the GBR increasing by 15% between 1998 and 2021. We conclude that GBR coral assemblages are systematically shifting away from past compositional states, a detectable trend on a decadal scale that is increasing across all GBR regions. Our results indicate that anomaly detection methods can contribute to both ecological research and management by providing a new way to quantify ecological novelty under climate change and anthropogenic pressures. Novelty can be estimated from compositional data without needing to establish reference baselines, providing a unique, non-equilibrium perspective that can advance ecological research aims and incorporate into conservation and restoration decision-making frameworks across different taxa and ecosystems. For reef assemblages, the widespread and consistent increase in novelty we identified echo warnings of a global ecological crisis. While a novel species assemblage does not necessarily translate into altered reef functionality or loss of resilience, our findings point to a new for reef management to account for the increasing gap between past reef states and emerging states of the future.

---

Intensifying human impacts (Steffen et al. 2015) are pushing abiotic envelopes away from the relative stability of the Holocene (Kerr et al. 2024, Summerhayes et al. 2024). While this is a whole-of-biosphere catastrophe, coral reefs are especially vulnerable to these anthropogenic impacts (Hughes et al. 2017). A cascade of human-induced stressors (Pandolfi et al. 2003, Mora et al. 2016) has led to reductions in coral cover (Emslie et al. 2024) and diversity (Gomez et al. 2018), shifts in assemblage structure (Hughes et al. 2018) and increases in the rate of community change (Blowes et al. 2019). As a consequence, we are observing the emergence of novel ecosystem states, driven by novel processes into configurations with no historical precedent (Jackson 2013, Kerr et al. 2024, Staples et al. 2025). This uniqueness in assemblage composition relative to the past is termed ecological novelty (Radeloff et al. 2015, Finsinger et al. 2017); novelty offers a way to quantify and measure ahistoric change in natural systems impacted by anthropogenic actions.

Novelty emerges in individual assemblages from altered abiotic and biotic conditions (Jackson 2013). These drivers of ecological novelty are not evenly distributed; in coral reefs, mass bleaching, predator outbreaks and water quality all differ substantially across space (Mellin et al. 2019). Ecological responses are similarly species-specific (Ortiz et al. 2018, McWilliam et al. 2020), creating winners and losers that push assemblages along alternate ecological pathways even where drivers of change are similar (Dornelas et al. 2019). The ultimate results are assemblages that may contain the same number and diversity, of organisms, as states of the past, even though actual species identities differ. Understanding and responding to anthropogenic threats both require distinguishing between different scenarios of assemblage change over time, such as between recovery and the emergence of a novel state, and between abrupt shifts (Johnstone et al. 2016), slow regime shifts (Hughes et al. 2013) and normal year-to-year compositional variation. Ecological novelty offers a quantitative tool with the potential to resolve these challenges.

Reliable measurement of novel ecological change remains a challenge. Novelty is a comparative measure of compositional uniqueness: therefore, the choice of an appropriate comparison is critical (Staples et al. 2025). Comparisons of ecological states over time, both in assemblage composition and summary variables like diversity, rely on defining a baseline “reference state” (McNellie et al. 2020, Czúcz et al. 2021). This reference state is assumed to be a climax state that an assemblage returns to following perturbation, often based on assemblage abundance or diversity (McNellie et al. 2020). However, identifying climax states, particularly in short time series of disturbance-prone ecosystems, is a challenge (Lindenmayer et al. 2012, Grainger et al. 2025) that will only intensify as anthropogenic impacts lead to press disturbances and potential community reassembly (Cooper et al. 2015, Rapacciuolo et al. 2019).

Ecological novelty is a form of anomaly detection, a field of research with methods that can distinguish outlying observations in complex, multidimensional data (Pimentel et al. 2014). While anomaly detection commonly uses one-class classification to distinguish inliers (non-novel) from outliers (novel), methods like Local Outlier Factor (LOF) provide a continuous estimate of an observation’s anomalousness, or novelty, relative to a set of neighbors in multidimensional space (Breunig et al. 2000). Notably, LOF uses a ratio of local neighborhood density rather than direct dissimilarities, relativizing novelty estimates to the compositional variability of the system. Applied to an ecological time series with past states as potential neighbors, LOF estimates novelty on a comparable scale across assemblages, without relying on pre-defined reference states. Novelty measured in this manner potentially fills a gap in the toolboxes of both researchers aiming to detect and understand ecological change, and managers attempting to monitor and conserve ecosystems under anthropogenic pressures (Reverter et al. 2022, Staples et al. 2025, Gonzalez-Rivero et al. 2025).

Here we test the ability of novelty measured via LOF to quantify ecological change through time at multiple spatial scales in coral reefs, as well as whether novelty can be reliably predicted by commonly measured ecological “condition” variables: coral cover, diversity, and year-to-year compositional change (turnover). We estimated LOF in 106 Great Barrier Reef time series collected by the Australian Institute of Marine Science Long-Term Monitoring Program and Marine Monitoring Program (Australian Institute of Marine Science 2014, 2015). These data extend from between 1993 to 2021 and contained coral cover and genus-growth form identifications. With novelty measured using LOF in these data, we asked the following questions:

1. Does novelty correlate with common ecological condition variables?,
2. Are reefs compositions on the Great Barrier Reef becoming novel, and if so, what is the trend in novelty over time?,
3. How do novelty trends differ among reefs and spatial regions across the GBR?, and
4. Can novelty estimated with LOF distinguish between common ecological trajectories, including (a) stable assemblage composition, (b) disturbance-recovery cycles, (c) abrupt change to a novel state and (d) gradual change in composition?

We hypothesized that novelty would be linked to the marked reductions in cover and diversity observed on the GBR (Hughes et al. 2018, 2019). We expected turnover to be positively correlated with novelty: novel states are likely to emerge from more rapid change than is normally observed in a system (Hobbs et al. 2014), and the GBR has been exposed to severe pulse disturbances that have driven rapid turnover (Hughes et al. 2018). We hypothesized that reef assemblages would become increasingly novel over the study period, in response to the increasing intensity and frequency of disturbance events (Emslie et al. 2024).

## Methods

All data processing and analyses were performed in R version 4.4.0 (R Core Team 2024).

Packages are listed in Section S1: Supplementary Methods.

We used coral assemblage time series from the Long-Term Monitoring Program run by the Australian Institute of Marine Science (AIMS) and the Marine Monitoring Program, a collaborative program between AIMS and the Great Barrier Reef Marine Park Authority for monitoring the condition of inshore reefs in the Great Barrier Reef Marine Park (Australian Institute of Marine Science 2014, 2015). These programs conduct benthic compositional surveys that estimate abundance via cover for a series of morphogenus/morphofamily categories that combine coral taxonomy and growth form. These categories include a broad set of scleractinian and soft corals, as well as octocoral *Heliopora* and the hydrozoan *Millepora*.

We excluded 71 reefs from the monitoring program with fewer than ten surveys and 143 surveys with potentially poor image resolution (where placeholder “non-*Acropora*” categories exceeded 10% of relative coral cover). Overall, data used here comprised 1,678 repeat surveys at 106 reefs along the GBR. Reef time series contained between ten and 27 surveys (median = 16). Excluding initial surveys, 1,060 of surveys were annual (63.17%), 503 (30.0%) were biennial, and nine were conducted after longer sampling gaps. Time series sampling commenced between 1993 and 1997 for 47 reefs, and 2005 and 2006 for 60 reefs following a change to monitoring sampling protocols.

### Deriving ecological variables

Each time series survey from each reef was treated as a point in multidimensional assemblage space. From this point, we estimated six ecological variables: coral cover, diversity, change in cover and diversity from the prior survey, temporal beta diversity (“turnover”) and Local Outlier Factor (“novelty”).

Coral cover was the summed cover of all coral categories in each survey. These cover estimates were used to calculate first-order differences, subtracting the prior year’s cover from each cover estimate, to reflect change in coral cover between surveys. All non-cover ecological variables were calculated using relative coral morphocategory cover that summed to one in each survey.

We estimated diversity as Hill number of order 1 (H1), equivalent to the exponent of Shannon’s Index. We also used first-order differencing on H1 estimates within each time series to reflect diversity change between surveys.

Turnover and novelty were estimated with Bray-Curtis dissimilarities using the square-root transformed relative abundance of coral morphocategories. Turnover was estimated as the Bray-Curtis dissimilarity between each survey and the prior year’s survey in the time series (Anderson et al. 2011), divided by the number of years since the last survey.

### Novelty measurement

Novelty was estimated using Local Outlier Factor (LOF), an anomaly detection method that uses a ratio of neighborhood densities based on dissimilarity values (Breunig et al. 2000). For each survey in each time series, we calculated LOF using the most similar five past surveys, identified by Bray-Curtis dissimilarity (full LOF specification is located as Section S1:

Supplementary Methods). Resulting LOF values were the ratio of survey-neighborhood dissimilarities divided by intra-neighborhood dissimilarities. An LOF ratio at or below one reflected a survey close to past assemblages, and thus low novelty value (Fig. 1A). High LOF values was an indication that the reef survey was outside the range of past observations. (Fig. 1B-C). Separate LOF values were calculated for all surveys in a time series excluding the first five (which had insufficient past neighbors for LOF calculation), for a total of 1,148 surveys.

**Figure 1:**
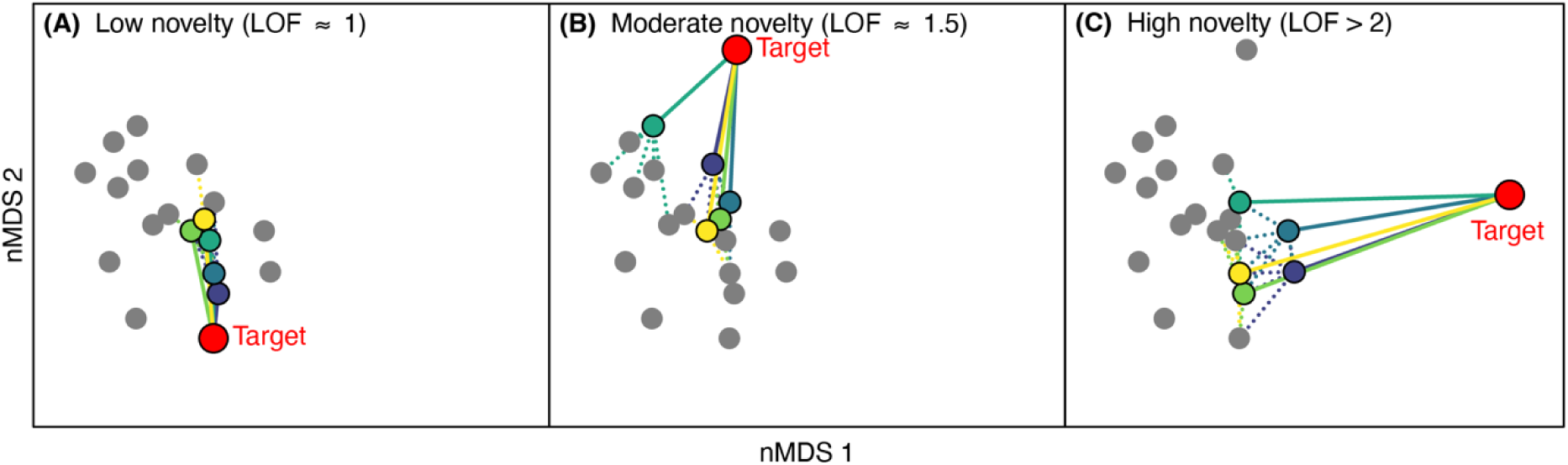
Examples of novelty estimated using Local Outlier Factor and hypothetical trends in novelty over time. (A) Low, (B) moderate and (C) high novelty in reef assemblage ordinations (2016-2018 surveys at Hastings Reef, Wet Tropics). Each point corresponds to a reef assemblage from a single year’s survey. Local Outlier Factor (LOF) estimates novelty as the ratio of two dissimilarity of an observation to its closest neighbors (n = 5, coloured points and lines) and the dissimilarity of each neighbor to their neighbors (dottes lines).

### Data analysis

#### Predictive novelty model

We used LOF novelty values (Box-Cox transformed with λ = −4.8. See Section S1: Supplementary Methods) as the response variable in a linear mixed-effects model. Model fixed effects included calculated ecological variables: coral cover, change in cover, diversity, change in diversity and turnover. We also added two fixed effects that reflected time series features that potentially skewed LOF estimation: time since last survey, a measure of the time lag between surveys, and time series length, the total survey count in each time series. Finally, we added survey year as a fixed effect to estimate any temporal trend in novelty over our survey period.

All fixed effects, with exception of survey year, were z-normalized to μ = 0 and σ = 1 to allow direct comparison of slopes as standardized effect sizes. Survey year was transformed by subtracting the first year in the dataset with LOF estimates, 1998, from survey year, and dividing the result by the standard deviation of survey year across the entire dataset. This scaled survey year values to σ = 1 so the survey year slope could be compared with other fixed effects while ensuring the model intercept estimated novelty from the start of the survey period instead of mean survey year.

We fit this model with four sets of random effects to reflect spatiotemporal data structuring. First, we fit random intercepts for survey year (n = 29) and inshore versus offshore reefs (n = 2). We also fit random intercepts, and random slopes with survey year, for each reef (n = 106) nested within random intercepts for latitudinal GBR regions (n = 5).

This model tested three hypotheses. First, slopes of other ecological variables with novelty tested how well these variables could predict novelty estimated by LOF. Second, the model intercept and survey year slope provided, respectively, an estimate of mean GBR-wide novelty at the start of the study period, and the trend in novelty over time. Finally, reef-level random intercepts and slopes described the variation of novelty among study reefs.

#### Reef and region-level novelty trends

Reef-level novelty trends were estimated using reef and region novelty random slopes and intercepts from the predictive model. These random effects described how much variation in novelty (random intercepts) and trend in novelty over time (random slopes) was clustered within reefs, and which reefs exhibited trends that differed from the GBR-wide estimates from the model fixed effects.

To examine regional trends in novelty over time, we extracted the fixed-effect residuals from our predictive model, excluding the effect of survey year. We used these residuals as the response variable in a mixed-effect generalized additive model. We fit survey year as a spline-based fixed effect with separate splines estimated for the five GBR regions and inshore and offshore reefs, with reef included as a random intercept. Additive models were used to detect non-linear novelty trends for regions over the study period to contrast the overall linear slope of novelty with survey year in our predictive model.

#### Diagnosing assemblage trajectories

We identified four primary ecological trajectories that may be detectable via novelty trends: (1) a stable composition, (2) disturbance followed by recovery of the pre-disturbance state, (3) an abrupt change to a new, novel, stable state and (5) gradual compositional movement away from past states. While these types of ecological change may combine or overlap in the same reef, we aimed to identify reefs that showcased each individual trend. We visualized novelty trend over time in these example reefs, alongside non-metric multidimensional scaling of reef assemblage composition, to test whether these distinct trajectories had a unique novelty signature.

#### Null models and sensitivity tests

Our measurement of LOF was potentially biased during early time series surveys, where the pool of past compositional neighbors was small, resulting in a sparse neighborhood that may have poor coverage of the assemblage’s compositional space. This meant that any increasing novelty trend over time would be conflated with a statistical sampling effect that increased novelty with the number of past time series observations. These were impossible to separate in the observed time series, as the number of past surveys was strongly correlated with survey year. While we could have removed early observations from each time series prior to modelling, reef time series were already short relative to reef recovery following disturbance (Johns et al. 2014). Instead, we isolated the sampling effect via null modelling, removing any temporal trend in reef composition with matrix permutation. We used the *permatfull* function in the vegan package (Oksanen et al. 2025), treating each 0.1% of morphocategory cover as an individual that could be randomly swapped among matrix cells. This process was unconstrained to remove potential temporal autocorrelation in taxon abundances. We repeated the permutation 999 times for each time series, each time remeasuring novelty for each survey (999 sets of 1,148 observations).

Novelty values in these null permutations retained the sampling effect of neighbor count as LOF was estimated sequentially along a time series, but removed any potential novelty trend over time.

We repeated the predictive model structure described above 999 times, each time substituting a set of null permuted novelty estimates as the response variable. We retained the observed values for all fixed and random effects. These null models were used to generate distributions of parameter estimates when temporal trend in coral compositional change was removed, with only sampling effects remaining.

As well as sampling effects, novelty measured via LOF potentially conflated two types of temporal trends: a directional change where later assemblage states differed from earlier states, and increasing assemblage variability over time. We conducted several sensitivity tests to distinguish between these, comparing (1) trends in novelty residuals from the predictive model, (2) time series simulations and (3) comparison between “past-only” and “whole-of-time series” estimation of novelty (S1: Supplementary Methods). These tests provided evidence our novelty estimates reflected compositional movement away from past assemblage states (S2: Supplementary Results, Fig. S1-3).

## Results

### Prediction of ecological novelty

Our predictive novelty model explained 40.9% of novelty variation with fixed effects (marginal R^2^), 79.4% including random effects (an additional 38.5%), leaving 20.6% as unexplained residuals. We found novelty increased over time across the GBR (Fig. 2). Novelty also correlated negatively with coral cover and diversity and positively with turnover, time series length and time since the prior survey (Fig. 2, S4).

**Figure 2:**
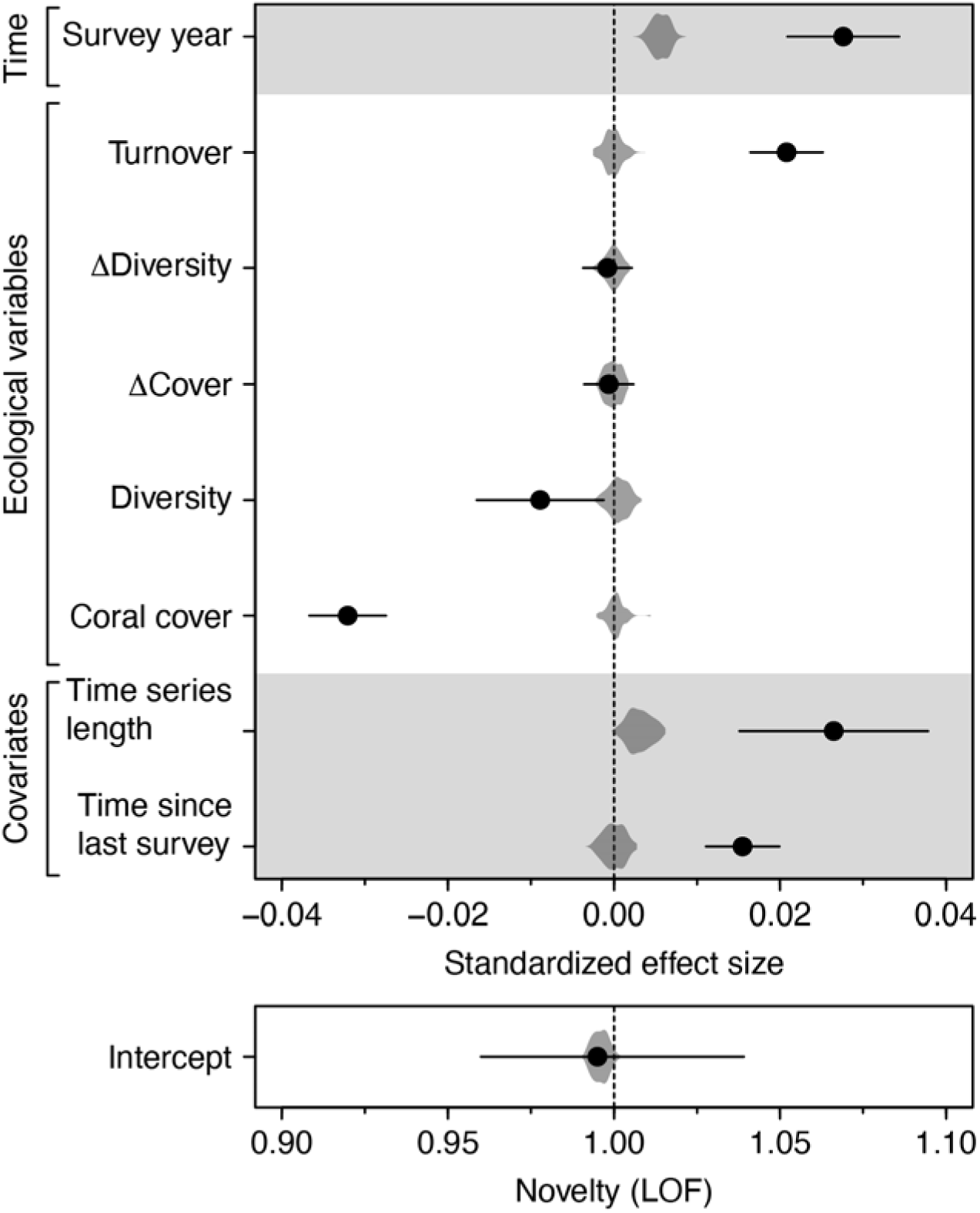
Coefficient plot of linear mixed-effects model predicting ecological novelty as a function of two covariates (time series length and time since last survey), five ecological variables and a linear trend through time (as survey year), including predicted novelty in 2000, an early, but well-sampled, survey year. Points are standardized effect sizes from observed data, with 95% confidence intervals. Density polygons represent the distribution of effect sizes from 999 null models with novelty calculated from randomly permuted compositional matrices. All fixed effects were z-normalized (μ = 0, σ = 1).

The model intercept was the predicted mean novelty across all surveyed reefs at the earliest survey year (1998), holding all other fixed effects at their GBR-wide mean. The intercept was 0.986, and not significantly different from one (Fig. 2, Table S1). This indicated that early time series surveys were not novel relative to prior assemblages. We also identified a positive relationship between novelty and survey year, indicating that novelty increased over time, with greater novelty in more recent surveys compared to earlier surveys (Fig. 2, Table S1). While the model slope was only linear in Box-Cox space, back-transformed to the ratio scale of LOF, predicted mean novelty increased from 0.986 in 1998 to 1.135 in 2021, an overall 15% increase. These positive novelty trends were detected while accounting for correlations between novelty other ecological variables (Fig. 2).

The null permuted data exhibited small positive slopes of novelty with survey year and time series length, pointing to a tendency for novelty to be higher in longer time series and with larger sets of past samples to select as LOF neighbors even in assemblages with no temporal autocorrelation (Fig. 2).

### Reef and region-level novelty trends

Reefs and regions contained systematically higher and lower novelty than the GBR-wide means, reflected in the spatially structured variation explained by model random effects (Table S1). At the regional scale, we observed increased novelty over time that was largely consistent with the linear GBR-wide trend (Fig. 3). Reefs in the northerly Cape York region were more novel than other regions across the entire study period, but especially between 2005 and 2012, and offshore reefs tended to be more novel than inshore reefs (Fig. 3). We identified 21 reefs (19.8%) with at least one survey with a novelty exceeding two, an indication of a highly novel assemblage, and an additional 25 reefs (23.58%) with a survey novelty between 1.5 and 2 (Table 1). Cape York, the most northerly GBR region, and Mackay-Whitsunday in the central GBR, contained the greater proportion of reefs with large novelty values, with seven of the eight Cape York reefs exhibiting a maximum novelty > 1.5 at some point since monitoring commenced (Table 1).

**Figure 3:**
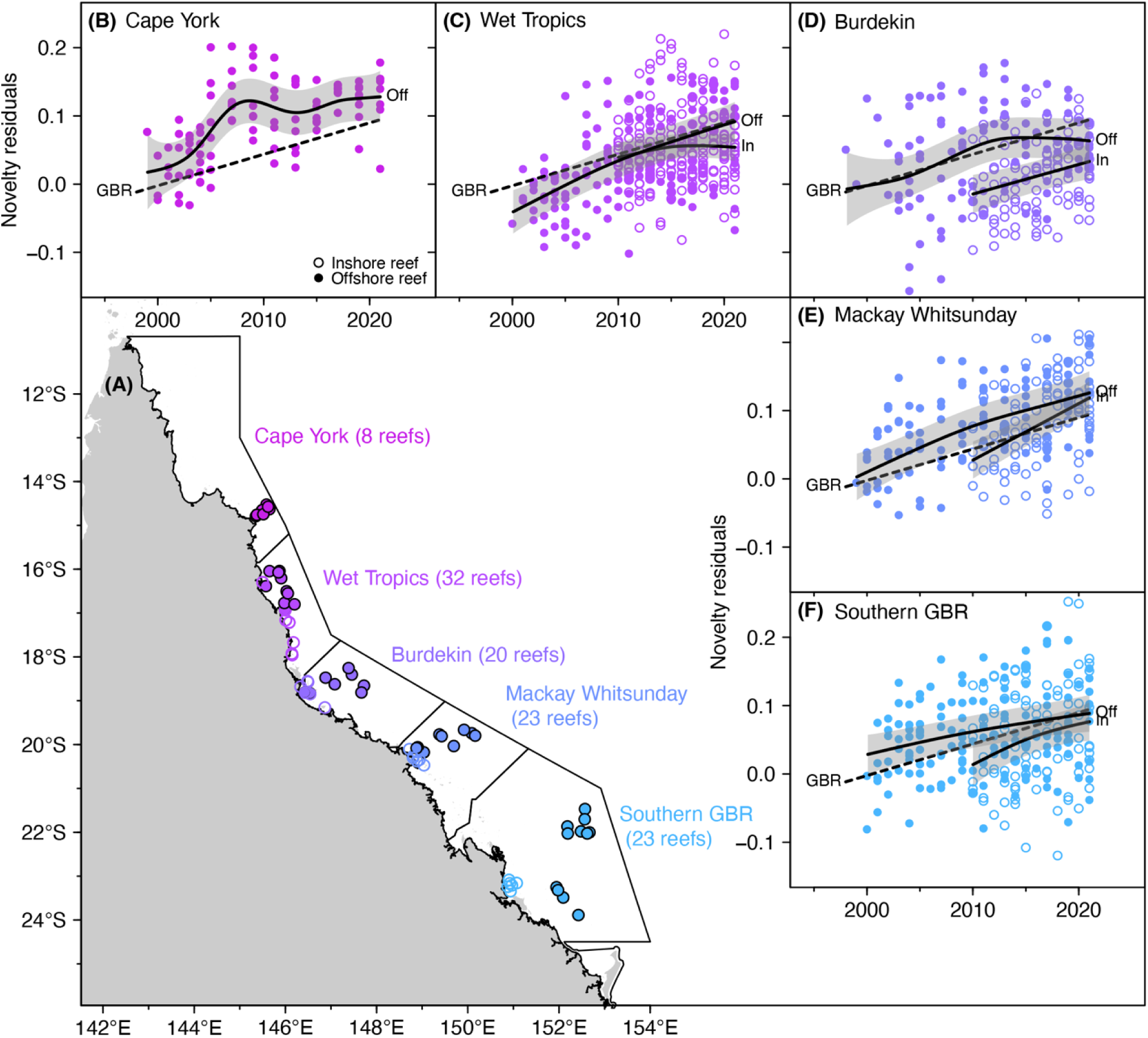
(A) Map of Great Barrier Reef Marine Park divided into spatial regions reflecting dominant catchment areas and reef sampling clusters. Numbers following names indicate number of reefs with time series compositional data. **(B-F)** Region-specific trends over time in novelty residuals, which were leftover novelty variation after accounting for all fixed effects in the predictive model except for time (survey year). Points are residual novelty observations. Dashed line (“GBR”) is the GBR-wide trend over time (time slope in Fig. 2). Solid lines and accompanying 95% confidence interval shading reflects region-specific trends in residual novelty over the study period in offshore (Off) and inshore (In) reefs.

**Table 1:** Reef count of the five spatial regions along the Great Barrier Reef (shown in. **Fig. 4A**), as well as number of reefs where the maximum observed novelty across all surveys exceeded two (highly novel), 1.5 (moderately novel) and below 1.5 (low or no novelty).

| Great Barrier Reef region | Reef count | Reef maximum novelty |  |  |
| --- | --- | --- | --- | --- |
| | | $x < 1.5$ | $1.5 \leq x < 2$ | $2 \leq x$ |
| Cape York | 8 | 1 (12.50%) | 4 (50.00%) | 3 (37.50%) |
| Wet Tropics | 32 | 20 (62.50%) | 7 (21.88%) | 5 (15.62%) |
| Burdekin | 20 | 17 (85.00%) | 3 (15.00%) | 0 (0.00%) |
| Mackay-Whitsunday | 23 | 10 (43.48%) | 4 (17.39%) | 9 (39.13%) |
| Southern GBR | 23 | 12 (52.17%) | 7 (30.43%) | 4 (17.39%) |

We added reef random intercepts and slopes to GBR-wide estimates (from Fig. 2) to predict reef-level novelty in 1998 and reef-level slope of in novelty over time. These represent linear expectations of novelty over time for a given reef holding all other predictors at the GBR-wide mean. As such, they are a coarse diagnostic for comparing among reefs rather than direct reef-level prediction of novelty over time. Observed reefs had intercepts and slopes well beyond null models, indicating novelty was spatially conserved within reefs even after accounting for model fixed effects (Fig. 4). Reef random intercepts and slopes were negatively correlated and represent a distribution of reef novelty trends (Fig. 4). The dominant axis described the timing of reef novelty. Reefs with large slopes and low intercepts exhibited lower novelty in early surveys and higher novelty in more recent surveys, whereas reefs with lower slopes and larger intercepts exhibited their highest novelty in earlier surveys. The orthogonal axis described the mean novelty observed across a reef’s surveys, with higher intercepts and slopes equating to more consistently high novelty, and vice versa (Fig. 4).

**Figure 4:**
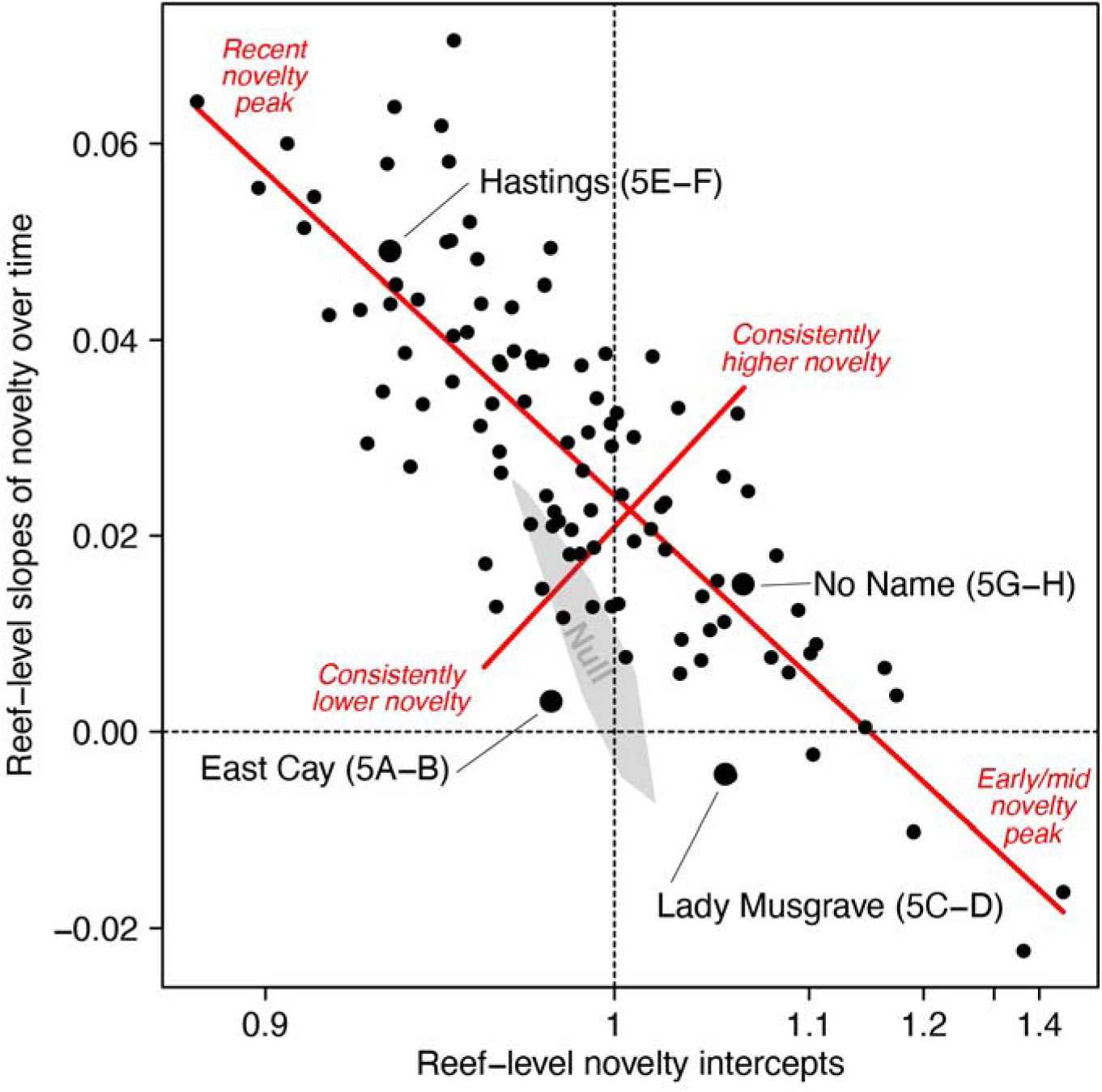
Reef-level novelty intercepts (x-axis, predicted novelty at mean survey year of 2013) and slope of novelty over time (y-axis). Each reef is represented by a point. The range of estimates from null permutation models enclosed in the convex hull, with the mean and 95% confidence intervals as the grey outlined point. Labelled reefs are examples of assemblage trajectories visualized in Figure 5, with subpanels in parentheses. Red lines reflect the primary and secondary axes of correlation between random intercepts and slopes, with accompanying description of their connection to reef-level novelty trends.

### Examples of reef trajectories

Novelty trends in individual reefs provided a useful description of four identified trajectories, which comprise the range of reef novelties observed from the model random effects (Fig. 4). East Cay, a remote reef in the southern GBR, exhibited a stable coral assemblage over the study period, which was reflected in novelty values consistently below 1.2 (Fig. 5A-B).

**Figure 5:**
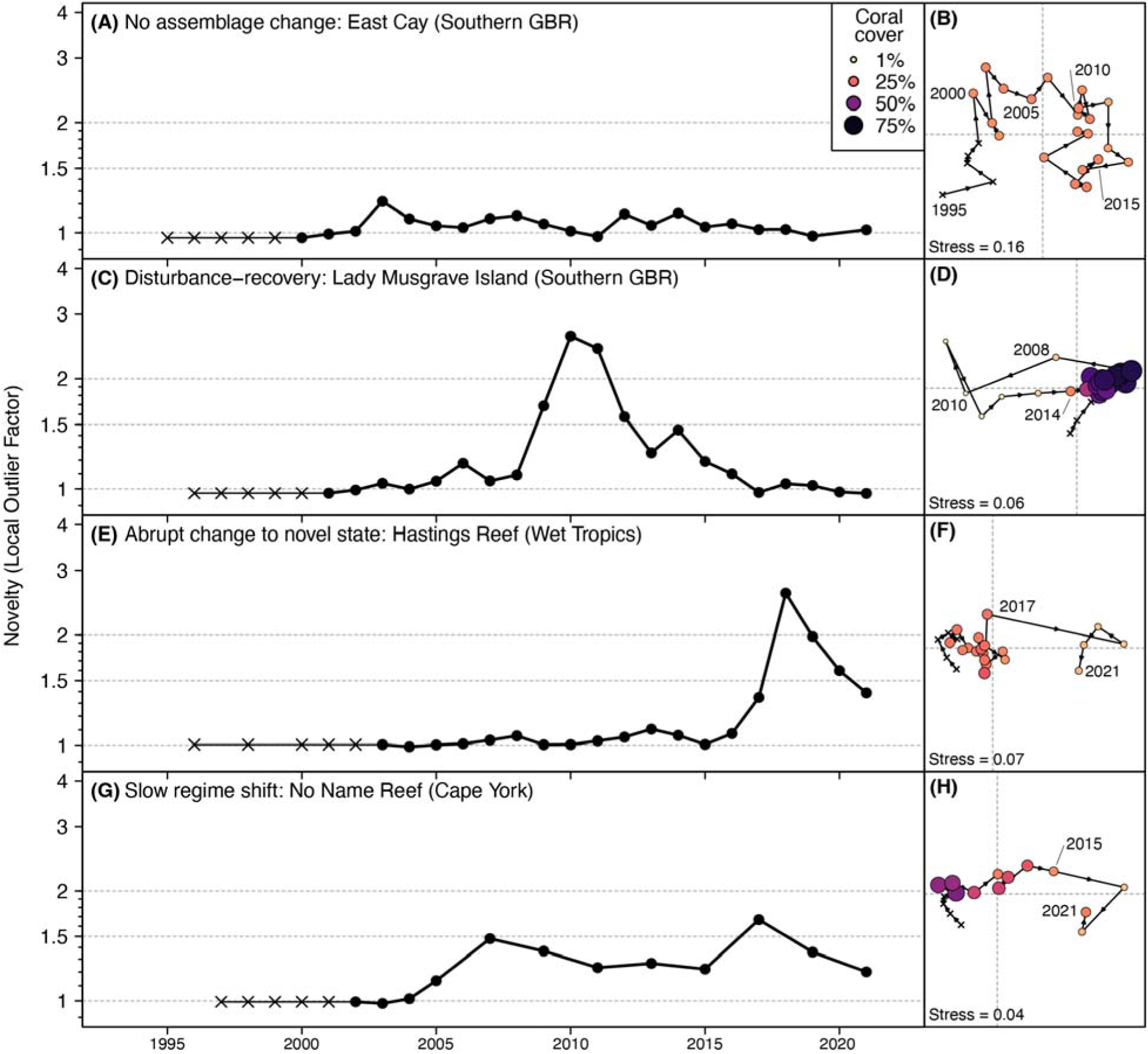
Example novelty over time for reefs representing four key scenarios: (A) stable assemblage over time, (C) disturbance followed by recovery, (E) abrupt change to a new stable state and (G) gradual directional change over time. Black crosses are initial five surveys in the time series where novelty was estimated. (B,D,F,H) Corresponding non-metric multidimensional scalings of reef composition based on Bray-Curtis dissimilarities, with surveys colored and sized based on total coral cover.

Lady Musgrave Island, in the Capricorn Bunker at the southern edge of the GBR, exhibited a classic disturbance-recovery cycle. In 2009, coral cover dropped to near zero following a disease outbreak and storm damage, but both cover and assemblage composition had recovered by 2015 (Fig. 5C-D). This trajectory manifested as a rapid increase in novelty followed by a decrease as the assemblage returned towards past states (Fig 5C-D).

Hastings Reef in the Wet Tropics suffered back-to-back mass bleaching events in 2016 and 2017. Novelty values peaked in 2018, the first survey following the second bleaching, but the assemblage did not return to its pre-disturbance state in subsequent surveys (Fig 5F). This potentially indicates the formation of a novel stable state. Novelty values decrease after the peak but remained elevated up until the final survey (Fig 5E). As the novel state stabilized it provided more past neighbors for Local Outlier Factor measurement, reducing novelty values (Fig 5E).

The final trajectory was defined by a gradual change in assemblage state. No Name Reef in the northern GBR exhibited a stable composition up until 2005 (Fig 5G-H). The reef assemblage began to change over time, impacted by several cyclones, multiple years of disease outbreaks followed by mass bleaching in 2016. Novelty values at No Name Reef were inflated but well below the values observed in previous abrupt change scenarios (Fig 5G-H). Instead, moderate novelties accumulated over time, at least up until 2015, as the assemblage moved away from early time series observations (Fig. 5H).

## Discussion

The Anthropocene portends a fundamental reassembly of ecological assemblages in response to changing abiotic and biotic regimes. Here we adopted an anomaly detection tool to estimate ecological novelty, the uniqueness of assemblage composition relative to the past. Over the last twenty years, we found that coral reef assemblages across the Great Barrier Reef are becoming increasingly novel, a trend that is increasing over time and in addition to volatility in coral cover (Emslie et al. 2024, Ceccarelli et al. 2026) and increasing rate of compositional change (Blowes et al. 2019). Our novelty trends do not, in isolation, implicate the drivers of directional ecological change or which taxa are changing (Staples et al. 2025), nor do they reflect altered reef function or loss of resilience. Our results explicitly point to a systematic shift in the coral composition of GBR reefs, showcasing how ecological novelty contributes to a multi-faceted description of assemblage trends over time. Coral reefs are not the only ecosystems responding to anthropogenic pressures (Blowes et al. 2019, Dornelas et al. 2019). Ecology is grappling with the need to detect both abrupt (Turner et al. 2020) and gradual (Hughes et al. 2013) compositional change as changes to disturbance regimes result in “long transient” systems (Francis et al. 2021) where equilibrium dynamics are challenging to detect (Rapacciuolo et al. 2019, Grainger et al. 2025). Our ability to understand and respond to future ecosystem dynamics relies on our ability to measure and compare important system attributes. Our novelty measurements using Local Outlier Factor (LOF) represent a powerful tool for ecological inference and management, converting local assemblage specifics into a comparable ratio scale that reflects compositional distance from past assemblage states.

### Situating ecological novelty among ecological summary variables

Our estimates of ecological novelty represent a change in assemblage composition compared to the past, but we hypothesized these changes would be accompanied by shifts in assemblage abundance (proxied by coral cover), diversity and the rate of year-to-year change. We found evidence for all three hypothetical correlations: higher novelty corresponded with lower cover and diversity and increased turnover. These relationships were only observed in the real-world reef time series and not in null permuted data. This suggests novelty is not mathematically dependent on other ecological variables and instead, novel reef states emerged alongside change in other ecological attributes. This is a common pattern, particularly when observing the impact of disturbance (Sousa 1984). Pulse disturbance removes individuals and vulnerable species, lowering cover and diversity and results in a different system from the pre-disturbance state, signalled by rapid turnover (Hidetoshi et al. 2022). These changes manifest as high novelty if the disturbed state had not been previously observed (e.g., in Lady Musgrave Island). Reef dynamics on the timescale of our data are largely driven by these pulse disturbances (Pandolfi 2002), and increases in the frequency and intensity of pulse disturbances are likely to be key drivers of future reef degradation (Mora et al. 2016, Emslie et al. 2024). We would expect novelty to correlate differently where, for instance, novelty is driven by biotic change like invasive species or in systems where pulse disturbances are rare (Harris et al. 2018). Even within our studied reefs, the spectrum of novelty trends over time reflects not only differing disturbance histories, but taxa-specific responses to disturbance (Johns et al. 2014). The life-history-based and time-lagged interaction between drivers and consequences of ecological change are challenging to unravel (Lalechère et al. 2025). By mapping onto hypothesized reef assemblage change, our results are evidence that novelty measures what it was designed to measure, while scaling very different reef assemblages onto a common ratio scale that allows for joint modelling and comparison.

Despite these correlations, 60% of variation in novelty was not explained by these more traditional ecological metrics. For example, we observed high reef novelty at times of high cover, high diversity and and low turnover (Fig. S1). Ecological change can occur without underlying changes to abundance and diversity (Hillebrand et al. 2018). Heavily invaded forests in Hawaii, for instance, have maintained ecosystem function and even increased in regional and local diversity over time (Mascaro et al. 2012). Press-like disturbances that increase slowly over time can exceed tipping points and create abrupt change (Andersen et al. 2009), but can also manifest as slow regime changes that are difficult to separate from background stochasticity and disturbance-recovery cycles (Hughes et al. 2013, Hidetoshi et al. 2022). Novelty has the potential to identify assemblage change that occurs without heightened turnover or diversity loss; our example reefs highlight how novelty maps reliably onto common assemblage trajectories, including both recovery from disturbance and the emergence of novel states.

Novelty as measured here is an ecological pattern that reflects altered coral composition (Staples et al. 2025). High novelty does not necessarily indicate whether reef functioning has changed, or even if novelty indicates a less desirable reef state from a conservation or climate resilience perspective. The value of novelty as measured here is the ability to decompose change in multidimensional assemblage compositions into a simple metric: how different the assemblage is to the past. This distinguishes novelty from other temporally comparative measures like turnover, or temporal beta diversity (Anderson et al. 2011). We applied a spatially restricted definition of novelty, whereby a reef state could be novel in one time series even if it had been previously observed in another place (as in Staples et al. 2022). The choice of reference set for comparison is key to understanding the ecological meaning of novelty (Staples et al. 2025). While novelty as a concept can be applied across space (e.g., Burke et al. 2019), our local-space and through-time characterisation grounded novelty in the past of each reef, allowing us to explore how local assemblage states have altered on a decadal scale.

Novelty measured by LOF offers three key features. First, LOF swaps the pre-defined description of a reference state for a data-driven specification of a reference set. By doing so, we let the data describe what is normal, and by extension, novel. This avoids the in-depth expertise needed to establish a reference state across a diverse set of reefs (e.g., Santavy et al. 2022), or proxying climax reef states using condition variables like coral cover (e.g., Guest et al. 2018). The second feature of LOF is that it measures novelty to a set of compositional neighbors, rather than a single neighbor (e.g., the most similar neighbor: Radeloff et al. 2015). This allowed us to avoid edge effects and identify novelty for multiple years even where nearest-neighbor distances were low (e.g., Lady Musgrave Island and Hastings Reef). The final feature is that LOF relativizes novelty using the density of the local compositional neighborhood. The ratio nature of LOF results in conservative estimates; to be novel, an observation must exist in a lower density area of compositional space than its most similar neighbors. This means that the same novelty value can emerge two ways: as a large dissimilarity from a low-density neighborhood, and as a small dissimilarity from a high-density neighborhood. Whether these are ecologically comparable will depend on local specifics, and that is where we feel ecological novelty is best situated. Not as a replacement for other ecological signals, but as a new perspective that complements existing summary statistics, improving our ability to compare and understand ecological change over time.

The relative nature of Local Outlier Factor, and ecological novelty generally, may act as a useful rate-based proxy for stability. Low novelty despite disturbance, like in East Cay (Fig. 5A-B) could indicate a system’s resistance to perturbation (Donohue et al. 2016). A rapid decline in novelty after an increase, as in Lady Musgrave Island (Fig. 5C-D) may reflect system resilience, a return to a non-novel past state (Holling 1973). Novelty trends could be used to estimate return time in compositional state, rather than benchmarks of cover or biomass, without needing to define a known climax state. Used in this way, novelty reflects the “stability of a rate”, rather than “stability of a state”; novelty is the rate of compositional change relative to the past, rather than the distance to a pre-defined climax state (Van Meerbeek et al. 2021).

### An increasingly novel Great Barrier Reef

Reef ecosystems are in crisis from increasingly frequent and intense disturbances (Emslie et al. 2024) driving reductions in population sizes (Hughes et al. 2018) and recruitment and recovery rates (Ortiz et al. 2018). We found a GBR-wide trend of increasing novelty, where reef assemblages have moved increasingly further away from observed states of the 1990-2000s. Our increasing novelty trend over time was modest, which is to be expected when averaged across over a hundred reefs spanning much of the GBR’s latitudinal gradient at a yearly grain. Estimates of zero-to-low novelty in 1998 was not an indication that reefs were undegraded. Anthropogenic impacts on reef systems pre-date our earliest surveys, with some human impacts commencing 150 years ago (McCulloch et al. 2003, Roff et al. 2013). In our study period, observed states with high coral cover and low novelty were still shifted baselines (Soga and Gaston 2018), representing “best-on-offer” reef states rather than a climax state absent human degradation (McNellie et al. 2020). Shifting baselines are a structural issue in temporal ecology that complicates any comparative measure (Gatti et al. 2015), and while LOF can mitigate these impacts by measuring novelty using multiple neighbors, relativized by neighborhood density, it does not eliminate them.

The lower predicted novelty in inshore reefs is evidence of how comparative measures, including novelty, are a function of time series context. Inshore reefs have been exposed to the cumulative impacts of land clearing over the past 150 years (Fabricius et al. 2005, Ceccarelli et al. 2020). In such cases, low novelty is not an encouraging sign, but rather a signal of a stable assemblage over the study period. Poor water quality has created strong abiotic filters to select for stress-tolerant coral species (Fabricius et al. 2005, DeVantier et al. 2006), locking reefs into marginal, resilience-focussed states (MacNeil et al. 2019). The ability to reliably measure the magnitude of assemblage change relative to the past comes with the loss of directionality: novelty collapses complex compositional change down to a single measure of “similar” versus “dissimilar”, which does not necessarily relate to whether the change is desirable.

Where ecological composition has changed, our ecological expectations may need to change too (Bellwood et al. 2019). A system with a novel composition may not contain the same taxa, support the same community members or provide the same levels, and types, of ecosystem functions as it did in the past (Graham et al. 2014). Reefs, for instance, may reassemble into states with historical levels of coral cover, and even diversity, with a new, novel assemblage (Graham et al. 2014). Novel systems weaken the ability for high cover, for instance, to proxy similarity to past states. Our results here highlight how novelty can provide unique information across multiple spatial scales that compliments existing ecological perspectives.

## Conclusions

In the Anthropocene, more than ever, we need to reliably measure assemblage-level ecological change. Ecological novelty in time series describes how far assemblages have shifted from past compositional states, but this is a pattern that does not implicate processes or ascribe value (e.g., “undegraded” versus “degraded”). Our results here highlight that novelty contains unique information that cannot be gleaned from a complex ecological dataset, like a time series, using a combination of other common metrics. We provide a promising first step to ground the measurement of ecological novelty in the real-world misfortune of reef systems. Most reefs along the Great Barrier Reef are moving away from historic states, concomitant with a period of increasing pressures imposed by climate change. While reef systems are particularly vulnerable to human impacts, the entire biosphere is being altered by human actions (Ellis 2011, Lenton et al. 2025). As human impacts intensify, we need new quantitative tools to describe the ecological consequences. Novelty estimated via anomaly detection methods offers a new perspective that can synergize with existing ecological variables to identify assemblage change over time, including the potential emergence of new compositional states, with untapped potential to inform ecological inquiry.

## Acknowledgements

This work was funded by the Australian Research Council’s (ARC) Discovery Early Career Research Award (DE240100398) awarded to TLS and the Great Barrier Reef Foundation Reef Trust Partnership, Stage 2, Critical Monitoring Program funding for an “Indicator framework for assessing the condition and recovery capacity of coral reef habitats in the Great Barrier Reef”. We gratefully acknowledge the work of the Australian Institute of Marine Science and the Great Barrier Reef Marine Park Authority for the development and maintenance of their long-term reef monitoring programs, which were used to produce this work. We thank A. Thompson for valuable suggestions that meaningfully improved our work here.

## Author contributions

The ideas for the paper were conceived by TLS and SWK. TLS conducted all analyses and developed analytical approaches with support from SWK. KC provided invaluable reef expertise and advice regarding the AIMS monitoring data. TLS wrote the paper with input from all other authors.

## Notes

https://doi.org/10.5281/zenodo.18332471

### Competing Interest Statement

The authors have declared no competing interest.

## Reference list

Andersen, T., J. Carstensen, E. Hernández-García, and C. M. Duarte. 2009. Ecological thresholds and regime shifts: approaches to identification. Trends in Ecology & Evolution 24:49–57.

Anderson, M. J., T. O. Crist, J. M. Chase, M. Vellend, B. D. Inouye, A. L. Freestone, N. J. Sanders, H. V. Cornell, L. S. Comita, K. F. Davies, S. P. Harrison, N. J. B. Kraft, J. C. Stegen, and N. G. Swenson. 2011. Navigating the multiple meanings of β diversity: a roadmap for the practicing ecologist. Ecology Letters 14:19–28.

Australian Institute of Marine Science. 2014. Great Barrier Reef Marine Monitoring Program - Coral (MMP).

Australian Institute of Marine Science. 2015. AIMS Long-term Monitoring Program.

Bellwood, D. R., M. S. Pratchett, T. H. Morrison, G. G. Gurney, T. P. Hughes, J. G. Álvarez-Romero, J. C. Day, R. Grantham, A. Grech, A. S. Hoey, G. P. Jones, J. M. Pandolfi, S. B. Tebbett, E. Techera, R. Weeks, and G. S. Cumming. 2019. Coral reef conservation in the Anthropocene: Confronting spatial mismatches and prioritizing functions. Biological Conservation 236:604–615.

Blowes, S. A., S. R. Supp, L. H. Antão, A. Bates, H. Bruelheide, J. M. Chase, F. Moyes, A. Magurran, B. McGill, I. H. Myers-Smith, M. Winter, A. D. Bjorkman, D. E. Bowler, J. E. K. Byrnes, A. Gonzalez, J. Hines, F. Isbell, H. P. Jones, L. M. Navarro, P. L. Thompson, M. Vellend, C. Waldock, and M. Dornelas. 2019. The geography of biodiversity change in marine and terrestrial assemblages. Science 366:339–345.

Breunig, M. M., H.-P. Kriegel, R. T. Ng, and J. Sander. 2000. LOF: Identifying Density-Based Local Outliers. SIGMOD Rec. 29:93–104.

Burke, K. D., J. W. Williams, S. Brewer, W. Finsinger, T. Giesecke, D. J. Lorenz, and A. Ordonez. 2019. Differing climatic mechanisms control transient and accumulated vegetation novelty in Europe and eastern North America. Philosophical Transactions of the Royal Society B: Biological Sciences 374:20190218–20190218.

Ceccarelli, D. M., M. J. Emslie, M. Logan, P. Bray, A. R. Campili, N. Cantin, A. Cole, M. J. Jonker, T. H. Sinclair Taylor, and B. Stephenson. 2026. Substantial impacts from the 2024 summer reverse five years of coral cover gains on the Great Barrier Reef. Coral Reefs.

Ceccarelli, D. M., R. D. Evans, M. Logan, P. Mantel, M. Puotinen, C. Petus, G. R. Russ, and D. H. Williamson. 2020. Long-term dynamics and drivers of coral and macroalgal cover on inshore reefs of the Great Barrier Reef Marine Park. Ecological Applications 30:e02008–e02008.

Cooper, J. K., M. Spencer, and J. F. Bruno. 2015. Stochastic dynamics of a warmer Great Barrier Reef. Ecology 96:1802–1811.

Czúcz, B., H. Keith, J. Maes, A. Driver, B. Jackson, E. Nicholson, M. Kiss, and C. Obst. 2021. Selection criteria for ecosystem condition indicators. Ecological Indicators 133:108376–108376.

DeVantier, L. M., G. De’ath, E. Turak, T. J. Done, and K. E. Fabricius. 2006. Species richness and community structure of reef-building corals on the nearshore Great Barrier Reef. Coral Reefs 25:329–340.

Donohue, I., H. Hillebrand, J. M. Montoya, O. L. Petchey, S. L. Pimm, M. S. Fowler, K. Healy, A. L. Jackson, M. Lurgi, D. McClean, N. E. O’Connor, E. J. O’Gorman, and Q. Yang. 2016. Navigating the complexity of ecological stability. Ecology Letters 19:1172–1185.

Dornelas, M., N. J. Gotelli, H. Shimadzu, F. Moyes, A. E. Magurran, and B. J. McGill. 2019. A balance of winners and losers in the Anthropocene. Ecology Letters 22:847–854.

Ellis, E. C. 2011. Anthropogenic transformation of the terrestrial biosphere. Philosophical Transactions of the Royal Society A: Mathematical, Physical and Engineering Sciences 369:1010–1035.

Emslie, M. J., M. Logan, P. Bray, D. M. Ceccarelli, A. J. Cheal, T. P. Hughes, K. A. Johns, M. J. Jonker, E. V. Kennedy, J. T. Kerry, C. Mellin, I. R. Miller, K. Osborne, M. Puotinen, T. Sinclair-Taylor, and H. Sweatman. 2024. Increasing disturbance frequency undermines coral reef recovery. Ecological Monographs 94:e1619–e1619.

Fabricius, K., G. De’ath, L. McCook, E. Turak, and D. McB. Williams. 2005. Changes in algal, coral and fish assemblages along water quality gradients on the inshore Great Barrier Reef. Marine Pollution Bulletin 51:384–398.

Finsinger, W., T. Giesecke, S. Brewer, and M. Leydet. 2017. Emergence patterns of novelty in European vegetation assemblages over the past 15 000 years. Ecology Letters 20:336–346.

Francis, T. B., K. C. Abbott, K. Cuddington, G. Gellner, A. Hastings, Y.-C. Lai, A. Morozov, S. Petrovskii, and M. L. Zeeman. 2021. Management implications of long transients in ecological systems. Nature Ecology & Evolution 5:285–294.

Gatti, G., C. N. Bianchi, V. Parravicini, A. Rovere, A. Peirano, M. Montefalcone, F. Massa, and C. Morri. 2015. Ecological Change, Sliding Baselines and the Importance of Historical Data: Lessons from Combing Observational and Quantitative Data on a Temperate Reef Over 70 Years. PLOS ONE 10:e0118581.

Gomez, C. G., A. Gonzalez, and H. M. Guzman. 2018. Multiscale change in reef coral species diversity and composition in the Tropical Eastern Pacific. Coral Reefs 37:105–120.

Gonzalez-Rivero, M., K. Crossman, A. Thompson, J. C. Ortiz, S. W. Kim, M. Emslie, A. S. Hoey, M. Hoogenboom, K. Bairos-Novak, E. McClure, J. M. Pandolfi, P. J. Mumby, M. Logan, B. Schaffelke, T. Staples, and K. E. Fabricius. 2025. Beyond Coral Cover A framework for assessing the condition of coral reef habitats and informing conservation targets. bioRxiv:2025.11.01.686047–2025.11.01.686047.

Graham, N. A. J., J. E. Cinner, A. V. Norström, and M. Nyström. 2014. Coral reefs as novel ecosystems: embracing new futures. Current Opinion in Environmental Sustainability 7:9–14.

Grainger, T. N., K. Stark, C. Song, M. A. Barbour, and R. M. Germain. 2025. The Equilibrium Conundrum. Ecology Letters 28:e70232–e70232.

Guest, J. R., P. J. Edmunds, R. D. Gates, I. B. Kuffner, A. J. Andersson, B. B. Barnes, I. Chollett, T. A. Courtney, R. Elahi, K. Gross, E. A. Lenz, S. Mitarai, P. J. Mumby, H. R. Nelson, B. A. Parker, H. M. Putnam, C. S. Rogers, and L. T. Toth. 2018. A framework for identifying and characterising coral reef “oases” against a backdrop of degradation. Journal of Applied Ecology 55:2865–2875.

Harris, R. M. B., L. J. Beaumont, T. R. Vance, C. R. Tozer, T. A. Remenyi, S. E. Perkins-Kirkpatrick, P. J. Mitchell, A. B. Nicotra, S. McGregor, N. R. Andrew, M. Letnic, M. R. Kearney, T. Wernberg, L. B. Hutley, L. E. Chambers, M.-S. Fletcher, M. R. Keatley, C. A. Woodward, G. Williamson, N. C. Duke, and D. M. J. S. Bowman. 2018. Biological responses to the press and pulse of climate trends and extreme events. Nature Climate Change 8:579–587.

Hidetoshi, I., M. Adam, R. Stephen, B. Angus, and S. Katriona. 2022. Pulse and Press Disturbances Have Different Effects on Transient Community Dynamics. The American Naturalist 200:571–583.

Hillebrand, H., B. Blasius, E. T. Borer, J. M. Chase, J. A. Downing, B. K. Eriksson, C. T. Filstrup, W. S. Harpole, D. Hodapp, S. Larsen, A. M. Lewandowska, E. W. Seabloom, D. B. Van de Waal, and A. B. Ryabov. 2018. Biodiversity change is uncoupled from species richness trends: Consequences for conservation and monitoring. Journal of Applied Ecology 55:169–184.

Hobbs, R. J., E. Higgs, C. M. Hall, P. Bridgewater, F. S. Chapin, E. C. Ellis, J. J. Ewel, L. M. Hallett, J. Harris, K. B. Hulvey, S. T. Jackson, P. L. Kennedy, C. Kueffer, L. Lach, T. C. Lantz, A. E. Lugo, J. Mascaro, S. D. Murphy, C. R. Nelson, M. P. Perring, D. M. Richardson, T. R. Seastedt, R. J. Standish, B. M. Starzomski, K. N. Suding, P. M. Tognetti, L. Yakob, and L. Yung. 2014. Managing the whole landscape: Historical, hybrid, and novel ecosystems. Frontiers in Ecology and the Environment 12:557–564.

Holling, C. S. 1973. Resilience and Stability of Ecological Systems. Annual Review of Ecology and Systematics 4:1–23.

Hughes, T. P., M. L. Barnes, D. R. Bellwood, J. E. Cinner, G. S. Cumming, J. B. C. Jackson, J. Kleypas, I. A. van de Leemput, J. M. Lough, T. H. Morrison, S. R. Palumbi, E. H. van Nes, and M. Scheffer. 2017. Coral reefs in the Anthropocene. Nature 546:82–90.

Hughes, T. P., J. T. Kerry, A. H. Baird, S. R. Connolly, T. J. Chase, A. Dietzel, T. Hill, A. S. Hoey, M. O. Hoogenboom, M. Jacobson, A. Kerswell, J. S. Madin, A. Mieog, A. S. Paley, M. S. Pratchett, G. Torda, and R. M. Woods. 2019. Global warming impairs stock–recruitment dynamics of corals. Nature 568:387–390.

Hughes, T. P., J. T. Kerry, A. H. Baird, S. R. Connolly, A. Dietzel, C. M. Eakin, S. F. Heron, A. S. Hoey, M. O. Hoogenboom, G. Liu, M. J. McWilliam, R. J. Pears, M. S. Pratchett, W. J. Skirving, J. S. Stella, and G. Torda. 2018. Global warming transforms coral reef assemblages. Nature 556:492–496.

Hughes, T. P., C. Linares, V. Dakos, I. A. van de Leemput, and E. H. van Nes. 2013. Living dangerously on borrowed time during slow, unrecognized regime shifts. Trends in Ecology & Evolution 28:149–155.

Jackson, S. T. 2013. Perspective: Ecological Novelty is not New. Pages 63–65 Novel Ecosystems: Intervening in the New Ecological World Order.

Johns, K. A., K. O. Osborne, and M. Logan. 2014. Contrasting rates of coral recovery and reassembly in coral communities on the Great Barrier Reef. Coral Reefs 33:553–563.

Johnstone, J. F., C. D. Allen, J. F. Franklin, L. E. Frelich, B. J. Harvey, P. E. Higuera, M. C. Mack, R. K. Meentemeyer, M. R. Metz, G. L. W. Perry, T. Schoennagel, and M. G. Turner. 2016. Changing disturbance regimes, ecological memory, and forest resilience. Frontiers in Ecology and the Environment 14:369–378.

Kerr, M. R., A. Ordonez, F. Riede, and J.-C. Svenning. 2024. A biogeographic–macroecological perspective on the rising novelty of the biosphere in the Anthropocene. Journal of Biogeography 51:575–587.

Lalechère, E., J. Lenoir, R. Marrec, F. Essl, I. Kühn, and T. Ergon. 2025. Assessing biodiversity trends in a quasi-permanent non-equilibrium state. Trends in Ecology & Evolution 40:949–959.

Lenton, T. M., M. Milkoreit, S. Willcock, J. F. Abrams, D. I. Armstrong McKay, J. E. Buxton, J. F. Donges, S. Loriani, N. Wunderling, F. Alkemade, M. Barrett, S. Constantino, T. Powell, S. R. Smith, C. A. Boulton, P. Pinho, H. Dijkstra, P. Pearce-Kelly, R. M. Roman-Cuesta, and D. Dennis, editors. 2025. The Global Tipping Points Report 2025. University of Exeter, Exeter, UK.

Lindenmayer, D. B., G. E. Likens, A. Andersen, D. Bowman, C. M. Bull, E. Burns, C. R. Dickman, A. A. Hoffmann, D. A. Keith, M. J. Liddell, A. J. Lowe, D. J. Metcalfe, S. R. Phinn, J. Russell-Smith, N. Thurgate, and G. M. Wardle. 2012. Value of long-term ecological studies. Austral Ecology 37:745–757.

MacNeil, M. A., C. Mellin, S. Matthews, N. H. Wolff, T. R. McClanahan, M. Devlin, C. Drovandi, K. Mengersen, and N. A. J. Graham. 2019. Water quality mediates resilience on the Great Barrier Reef. Nature Ecology & Evolution 3:620–627.

Mascaro, J., R. F. Hughes, and S. A. Schnitzer. 2012. Novel forests maintain ecosystem processes after the decline of native tree species. Ecological Monographs 82:221–228.

McCulloch, M., S. Fallon, T. Wyndham, E. Hendy, J. Lough, and D. Barnes. 2003. Coral record of increased sediment flux to the inner Great Barrier Reef since European settlement. Nature 421:727–730.

McNellie, M. J., I. Oliver, J. Dorrough, S. Ferrier, G. Newell, and P. Gibbons. 2020. Reference state and benchmark concepts for better biodiversity conservation in contemporary ecosystems. Global Change Biology 26:6702–6714.

McWilliam, M., M. S. Pratchett, M. O. Hoogenboom, and T. P. Hughes. 2020. Deficits in functional trait diversity following recovery on coral reefs. Proceedings of the Royal Society B: Biological Sciences 287:20192628–20192628.

Mellin, C., S. Matthews, K. R. N. Anthony, S. C. Brown, M. J. Caley, K. A. Johns, K. Osborne, M. Puotinen, A. Thompson, N. H. Wolff, D. A. Fordham, and M. A. MacNeil. 2019. Spatial resilience of the Great Barrier Reef under cumulative disturbance impacts. Global Change Biology 25:2431–2445.

Mora, C., N. A. J. Graham, and M. Nyström. 2016. Ecological limitations to the resilience of coral reefs. Coral Reefs 35:1271–1280.

Oksanen, J., G. L. Simpson, F. G. Blanchet, R. Kindt, P. Legendre, P. R. Minchin, R. B. O’Hara, P. Solymos, M. H. H. Stevens, E. Szoecs, H. Wagner, M. Barbour, M. Bedward, B. Bolker, D. Borcard, T. Borman, G. Carvalho, M. Chirico, M. De Caceres, S. Durand, H. B. A. Evangelista, R. FitzJohn, M. Friendly, B. Furneaux, G. Hannigan, M. O. Hill, L. Lahti, C. Martino, D. McGlinn, M.-H. Ouellette, E. Ribeiro Cunha, T. Smith, A. Stier, C. J. F. Ter Braak, and J. Weedon. 2025. vegan: Community Ecology Package.

Ortiz, J.-C., N. H. Wolff, K. R. N. Anthony, M. Devlin, S. Lewis, and P. J. Mumby. 2018. Impaired recovery of the Great Barrier Reef under cumulative stress. Science Advances 4:eaar6127–eaar6127.

Pandolfi, J. M. 2002. Coral community dynamics at multiple scales. Coral Reefs 21:13–23.

Pandolfi, J. M., R. H. Bradbury, E. Sala, T. P. Hughes, K. A. Bjorndal, R. G. Cooke, D. McArdle, L. McClenachan, M. J. H. Newman, G. Paredes, R. R. Warner, and J. B. C. Jackson. 2003. Global trajectories of the long-term decline of coral reef ecosystems. Science 301:955–958.

Pimentel, M. A. F., D. A. Clifton, L. Clifton, and L. Tarassenko. 2014. A review of novelty detection. Signal Processing 99:215–249.

R Core Team. 2024. R: A Language and Environment for Statistical Computing.

Radeloff, V. C., J. W. Williams, B. L. Bateman, K. D. Burke, S. K. Carter, E. S. Childress, K. J. Cromwell, C. Gratton, A. O. Hasley, B. M. Kraemer, A. W. Latzka, E. Marin-Spiotta, C. D. Meine, S. E. Munoz, T. M. Neeson, A. M. Pidgeon, A. R. Rissman, R. J. Rivera, L. M. Szymanski, and J. Usinowicz. 2015. The rise of novelty in ecosystems. Ecological Applications 25:2051–2068.

Rapacciuolo, G., A. J. Rominger, N. Morueta-Holme, and J. L. Blois. 2019. Editorial: Ecological Non-equilibrium in the Anthropocene. Frontiers in Ecology and Evolution Volume 7–2019.

Reverter, M., S. B. Helber, S. Rohde, J. M. de Goeij, and P. J. Schupp. 2022. Coral reef benthic community changes in the Anthropocene: Biogeographic heterogeneity, overlooked configurations, and methodology. Global Change Biology 28:1956–1971.

Roff, G., T. R. Clark, C. E. Reymond, J. Zhao, Y. Feng, L. J. McCook, T. J. Done, and J. M. Pandolfi. 2013. Palaeoecological evidence of a historical collapse of corals at Pelorus Island, inshore Great Barrier Reef, following European settlement. Proceedings of the Royal Society B: Biological Sciences 280:20122100–20122100.

Santavy, D. L., S. K. Jackson, B. Jessup, J. Gerritsen, C. Rogers, W. S. Fisher, E. Weil, A. Szmant, D. Cuevas-Miranda, B. K. Walker, C. Jeffrey, P. Bradley, D. Ballantine, L. Roberson, H. Ruiz-Torres, B. Todd, T. Smith, R. Clark, E. Diaz, J. Bauzá-Ortega, C. Horstmann, and S. Raimondo. 2022. A biological condition gradient for coral reefs in the US Caribbean Territories: Part I. Coral narrative rules. Ecological Indicators 138:108805–108805.

Soga, M., and K. J. Gaston. 2018. Shifting baseline syndrome: causes, consequences, and implications. Frontiers in Ecology and the Environment 16:222–230.

Sousa, W. P. 1984. The Role of Disturbance in Natural Communities. Annual Review of Ecology and Systematics 15:353–391.

Staples, T. L., J. Blois, K. L. Cramer, E. T. Cunningham, M. Dornelas, S. G. Haberle, T. Heger, W. Kiessling, A. E. Magurran, A. O’Dea, A. M. Penny, V. C. Radeloff, J. A. Smith, W. Thuiller, J. W. Williams, and J. M. Pandolfi. 2025. A Conceptual Framework for Measuring Ecological Novelty. Global Ecology and Biogeography 34:e70005–e70005.

Staples, T. L., W. Kiessling, and J. M. Pandolfi. 2022. Emergence patterns of locally novel plant communities driven by past climate change and modern anthropogenic impacts. Ecology Letters 25:1497–1509.

Steffen, W., W. Broadgate, L. Deutsch, O. Gaffney, and C. Ludwig. 2015. The trajectory of the Anthropocene: The Great Acceleration. The Anthropocene Review 2:81–98.

Summerhayes, C. P., J. Zalasiewicz, M. J. Head, J. Syvitski, A. D. Barnosky, A. Cearreta, B. Fiałkiewicz-Kozieł, J. Grinevald, R. Leinfelder, F. M. G. McCarthy, J. R. McNeill, Y. Saito, M. Wagreich, C. N. Waters, M. Williams, and J. Zinke. 2024. The future extent of the Anthropocene epoch: A synthesis. Global and Planetary Change 242:104568–104568.

Turner, M. G., W. J. Calder, G. S. Cumming, T. P. Hughes, A. Jentsch, S. L. LaDeau, T. M. Lenton, B. N. Shuman, M. R. Turetsky, Z. Ratajczak, J. W. Williams, A. P. Williams, and S. R. Carpenter. 2020. Climate change, ecosystems and abrupt change: science priorities. Philosophical Transactions of the Royal Society B: Biological Sciences 375:20190105–20190105.

Van Meerbeek, K., T. Jucker, and J.-C. Svenning. 2021. Unifying the concepts of stability and resilience in ecology. Journal of Ecology 109:3114–3132.

