## Supplementary for "Anomaly detection uncovers emerging novelty in coral assemblages of the Great Barrier Reef"

**Supplementary material for**
Anomaly detection uncovers emerging novelty in coral assemblages of the Great Barrier Reef

**Authors:** Timothy L Staples^1*^, Sun Wook Kim^1^, Kerryn Crossman^2^, John M Pandolfi^1^

**S1: Supplementary methods**

R packages

We used the following R packages in our data processing and analyses: “DHARMa” (Hartig, 2020), "gamm4" (Wood & Scheipl, 2020), "gisaimsr" (Barneche & Logan, 2025), “lme4” (Bates et al., 2015), "MASS" (Venables & Ripley, 2002), “performance” (Lüdecke et al., 2021), "shape" (Soetaert, 2024), "sf" (Pebesma & Bivand, 2023), "tidyr" (Wickham et al., 2024), “usdm” , “vegan” (Oksanen et al., 2025) and “viridisLite” (Garnier et al., 2023).

Local Outlier Factor calculation

LOF calculation requires referring to a target survey (T), the target’s closest compositional neighbors (first-order neighbors or N_1_) and the closest neighbors of the target’s neighbors (second-order neighbors or N_2_). Equations 1 and 2 below are applied to both T and N_1_, and refer to S1 and S2 as generic survey references. Equation 3 is only applied to T relative to N_1_ and refers to them explicitly. LOF requires specifying the number of neighbors used for comparison, referred to as k.

LOF was calculated as follows (Sugiyama, 2016):

1. N_1_ observations were identified as the k-closest neighbours to T in compositional space (via the smallest Bray-Curtis dissimilarities). The dissimilarity (D) between T and the k-th N_1_ is called K-dissimilarity (D_k_).
2. For each N_1_, sets of N_2_ were identified as each N_1_’s k-closest neighbours. The number of k-closest neighbours is identical for T and N_1_.
3. Local Reachability Density (LRD) was calculated for T. LRD is a measure of density and estimates how densely T is surrounded by the set of N_1_ (eq. 2) using reachability distance (RD, eq. 1). Here T = S1 and N_1_ = S2. This equation is the inverse of the mean T to N_1_ RD, such that a high LRD results where T and N_1_ form a dense cluster (i.e., high density) and low LRD results where T and N_1_ sparsely populate compositional space (i.e., low density).
4. LRD is then calculated for each N_1_ using the sets of N_2_ (each N_1_ = S1 and corresponding N_2_ = S2). All LRDs for N_1_ used T’s K-dissimilarity for the reachability distance calculation.
5. Finally, LOF is calculated as the mean of LRD_T_-LRD_N1_ ratios.

| Eq. 1 (Reachability distance) | ${RD}_{k}\left( S1, S2 \right)=\max\left( D_{k}\left( S1 \right), D\left( S1,S2 \right) \right)$ |
| --- | --- |
| Eq. 2 (Local reachability density) | ${LRD}_{k}\left( S1 \right)= \left( \frac{1}{k}\sum_{i=1}^{k} {RD}_{k}\left( {S2}_{i}, S1 \right) \right)^{-1}$ |

| Eq. 3 (Local outlier factor) | $Local Outlier Factor {LOF}_{k}\left( T \right)= \frac{\frac{1}{k}\sum_{i=1}^{k} {LRD}_{k}(N1)}{{LRD}_{k}(T)}$ |
| --- | --- |

The resulting LOF values are ratios of T-N_1_ to N_1_-N_2_ neighbourhood densities. LOF values greater than one indicate a survey is anomalous, as it is in low density compositional space relative to its closest neighbors, on a ratio scale (e.g., LOF = 2 indicates the survey’s neighborhood is half as dense as its closest neighbors).

LOF neighbour choice

Local Outlier Factor requires an arbitrary choice of the number of neighbours. Here we set the neighborhood size to five, based on both statistical and ecological grounds. From a statistical perspective, using five neighbors avoids stochasticity from small neighborhoods while preserving the ability to detect novelty in the earlier surveys of reef time series. Ecologically, five years is roughly half of the estimated time a reef takes to recover following disturbance (Johns et al., 2014), providing a meaningful window of comparison to identify partially recovered states, or those recovering into novel compositions.

Other novelty scenarios may require different parameterization. In ideal datasets, we recommend the setting of ecologically informed neighborhood size, such as the expected recovery time from disturbance or the length of time organisms could persist in non-optimal conditions. Transparent and explicit consideration of novelty will be invaluable to assessing the limits of the measurement’s power to address unanswered ecological questions.

We validated our neighbourhood choice by estimating the correlation of LOF values using different neighborhood windows (Table S4). A neighborhood of k = 5 provided strong correlation to novelty values estimated with smaller (k = 3) and larger (k = 10) neighborhoods, acting as a best-case trade-off to use in main text analyses.

LOF/anomaly transformation for modelling

Local Outlier Factors are ratios, which present difficulties for satisfying model assumptions as a response variable. We used a Box-Cox transformation to linearize LOF values for modelling, testing λ scaling values from -10 to 10 in units of 0.1 using the boxcox function of the MASS package (Venables & Ripley, 2002). Log-likelihood was maximized at λ = -4.8 (Fig. S5), which we used as the transformation in main text analyses.

**S2: Supplementary results**

**Distinguishing between stochastic novelty and directional change in LOF measurements**

The Local Outlier Factor method is most used in a static multi-dimensional dataset, where all observations are potential neighbours. In these data, only observations that are outlying from most of the dataset will have high LOF values. However, in the time-forward nature of ecological inference and monitoring, we are often concerned with comparing the present to the past. We applied this framing to our measurement of LOF in this study, by measuring the anomaly of each survey only relative to past surveys. This meant that LOF values could be larger than they would be using the entire time series as potential neighbours, if a survey was anomalous relative to past surveys but not anomalous relative to future surveys. This brings LOF closer to our intention of measuring compositional change relative to the past. However, we risk conflating high LOF from increased compositional variance with high LOF from directional assemblage movement over time.

To parse these effects and test LOF's ability to measure past-looking compositional anomaly, we looked at three factors.

First, we examined the residual variance from our main-text predictive model. Increased LOF residual variance over time, particularly in top quantiles (e.g., 90%), would be an indication that main text trends may have been driven by increased compositional stochasticity, rather than a consistent trend away from past surveys. We found that the mean and quantiles of residuals were stable over time, an indication that LOF values were not systemically becoming more variable over the time series (Fig. S1).

Second, we ran two simulations, creating 999 time series of 24 years that only moved directionally over time (Fig. S2B) and only increased in variance over time (Fig. S2C). We overlaid a time-only slope of the observed GBR reef surveys (not including other predictors and covariates). We found that increasing compositional variance did not lead to a marked increase in LOF over time. This is because increased variance leads to not only an increase in the target-N1 dissimilarity, but also creates a less dense compositional space, where N1-N2 dissimilarities are inflated (see LOF equations above). This reduces LOF values as a ratio, even as dissimilarities increase. We found that directional change did increase LOF over time, evidence our past-only neighbourhood specification of LOF could capture this ecological trajectory (Fig. S2A). The observed GBR surveys had a stronger slope than even our directional change simulation, with a lower intercept. This suggests GBR reefs were more stable than our "linear change" scenario in the initial years, with accelerating directional change over time, which aligns with our major main-text conclusions.

The final test was to compare the "past-only" LOF specification of our main text results with a "whole-of-time series" LOF. In these data, surveys with high LOF values for both "past-only" and "whole-of-time series" are classic anomalies, assemblages that our outlying in compositional space. Surveys with a high "past-only" LOF but a lower "whole-of-time series" LOF contain directional information - they are outlying when the future is not considered, but not outlying when it is, suggesting they are part of a chain of transitional assemblage states.

The two sets of LOF values were highly correlated (Pearson's R of Box-Cox transformed LOF values = 0.809). Using only the 258 surveys with at least one LOF > 1.2, to exclude inlying surveys, we found 198 (76.74%) had "past-only" LOFs that were greater than their "whole-of-time series" LOFs, an indication of directional compositional change (Fig. S3). Another 46 had the same LOFs, 36 of which were the final survey of the time series (where neighbours in past-only and whole-of-time series were identical), and only 14 surveys had a lower "past-only" LOF (Fig. S3). The cluster of large LOF values (>2) that appear near-identical on the Box-Cox scale comprise 19 surveys, 13 of which had a larger "Past-only" LOF and six which were the same.

Larger "Past-only" LOFs occur where a survey is novel relative to the past, only to be less anomalous once future surveys have been filled in. This is the pattern we would expect where a time series is moving directionally through compositional space (as per Fig. S2B). The magnitude of difference between the two LOFs may be an indication of the stability of the survey, with a large difference indicating that the survey becomes much less novel once future observations are included, as we might expect if an abrupt change resulted in a new, novel, stable state.

A further exploration of the implementation of LOF to ecology, beyond our introduction here, is warranted. Such further work should examine how neighbourhood specification can impact novelty estimates, and how the inclusion of future information can impact interpretation. For our research aims, which include the potential for LOF to act as a monitoring tool, the past-to-present specification of LOF neighbourhood is an important feature. Our suite of supplementary results here provide evidence to support our main-text conclusions, that coral assemblages on the GBR are becoming increasingly novel.

**Supplementary references**

Barneche, D., & Logan, M. (2025). *gisaimsr: Assortment of GBR GIS Files*. https://github.com/open-AIMS/gisaimsr

Bates, D., Mächler, M., Bolker, B. M., & Walker, S. C. (2015). Fitting linear mixed-effects models using lme4. *Journal of Statistical Software*, *67*(1), 1–48. https://doi.org/10.18637/jss.v067.i01

Garnier, Simon, Ross, Noam, Rudis, Robert, Camargo, Pedro, A., Sciaini, Marco, Scherer, & Cédric. (2023). *viridis(Lite) - Colorblind-Friendly Color Maps for R*. https://doi.org/10.5281/zenodo.4678327

Hartig, F. (2020). DHARMa: Residual Diagnostics for Hierarchical (Multi-Level / Mixed) Regression Models. R package version 0.3.3.0. In *https://CRAN.R-project.org/package=DHARMa*. https://cran.r-project.org/package=DHARMa

Johns, K. A., Osborne, K. O., & Logan, M. (2014). Contrasting rates of coral recovery and reassembly in coral communities on the Great Barrier Reef. *Coral Reefs*, *33*(3), 553–563. https://doi.org/10.1007/s00338-014-1148-z

Lüdecke, D., Ben-Shachar, M. S., Patil, I., Waggoner, P., & Makowski, D. (2021). performance: An R Package for Assessment, Comparison and Testing of Statistical Models. *Journal of Open Source Software*, *6*(60), 3139. https://doi.org/10.21105/joss.03139

Oksanen, J., Simpson, G. L., Blanchet, F. G., Kindt, R., Legendre, P., Minchin, P. R., O’Hara, R. B., Solymos, P., Stevens, M. H. H., Szoecs, E., Wagner, H., Barbour, M., Bedward, M., Bolker, B., Borcard, D., Borman, T., Carvalho, G., Chirico, M., De Caceres, M., … Weedon, J. (2025). *vegan: Community Ecology Package*. https://CRAN.R-project.org/package=vegan

Pebesma, E., & Bivand, R. (2023). *Spatial Data Science: With applications in R*. Chapman and Hall/CRC. https://doi.org/10.1201/9780429459016

Soetaert, K. (2024). *shape: Functions for Plotting Graphical Shapes, Colors*. https://CRAN.R-project.org/package=shape

Sugiyama, M. (2016). Chapter 38 - Outlier Detection. In M. Sugiyama (Ed.), *Introduction to Statistical Machine Learning* (pp. 457–468). Morgan Kaufmann. https://doi.org/https://doi.org/10.1016/B978-0-12-802121-7.00049-2

Venables, W. N., & Ripley, B. D. (2002). *Modern Applied Statistics with S* (Fourth). Springer. https://www.stats.ox.ac.uk/pub/MASS4/

Wickham, H., Vaughan, D., & Girlich, M. (2024). *tidyr: Tidy Messy Data*. https://CRAN.R-project.org/package=tidyr

Wood, S., & Scheipl, F. (2020). *gamm4: Generalized additive mixed models using mgcv and lme4*. https://cran.r-project.org/package=gamm4

**Supplementary tables**

***Table S1:*** *Model summary table of predictive anomaly model. Dispersion and Uniformity tests were performed by residual simulation within the DHARMa package. Temporal autocorrelation test is the summary of Durbin-Watson tests performed across model residuals striated by site coordinates. Spatial autocorrelation test is the summary of Moran's I tests straited by survey year. While uniformity test was significant, test statistic was small.*

| **Fixed effects** | **Observed model** | | | **Null model estimate** | | |
| --- | --- | --- | --- | --- | --- | --- |
| Variable | Estimate | Std Error | *t*-value | Mean | 2.5% quantile | 97.5% quantile |
| Intercept | 0.0576 | 0.0187 | 3.0748 | 8.37e-3 | 6.94e-3 | 1.05e-2 |
| Time between survey years | 0.0155 | 0.0023 | 6.8034 | 3.83e-6 | -2.20e-3 | 2.39e-3 |
| Time series length | 0.0265 | 0.0058 | 4.5551 | 3.18e-3 | 4.57e-4 | 5.81e-3 |
| Coral cover | -0.0321 | 0.0024 | -13.5419 | 2.42e-4 | -1.78e-3 | 1.91e-3 |
| Diversity (Hill order 1) | -0.0089 | 0.0039 | -2.2704 | 4.72e-4 | -2.10e-3 | 3.27e-3 |
| dCover | -0.0006 | 0.0015 | -0.4093 | -1.60e-4 | -1.69e-3 | 1.54e-3 |
| dDiversity | 0.0008 | 0.0015 | 0.5249 | -8.53e-5 | -2.15e-3 | 1.70e-3 |
| Turnover | 0.0208 | 0.0022 | 9.2728 | -8.99e-5 | -2.11e-3 | 2.22e-3 |
| Survey year (time) | 0.0276 | 0.0034 | 8.0082 | 5.54e-3 | 3.45e-3 | 7.51e-3 |
| **Random effects** | Observed model | | Null model variance | | |  |
| Group | Variance | Std Dev | Mean | 2.5% quantile | 97.5% quantile |  |
| Reef:Region (Intercept) | 1.590e-3 | 0.0399 | 2.62e-5 | 0 | 6.14e-5 |  |
| Reef:Region (Slopes with survey year) | 5.595e-4 | 0.0237 | 1.04e-5 | 8.41e-8 | 4.02e-5 |  |
| Survey year | 3.969e-5 | 0.0063 | 8.20e-17 | 0 | 1.42e-5 |  |
| Region | 6.512e-4 | 0.0255 | 0 | 0 | 1.14e-5 |  |
| Inshore vs offshore | 3.912e-4 | 0.0198 | 5.61e-6 | 0 | 6.40e-5 |  |
| Residual | 1.734e-3 | 0.0416 | 8.69e-4 | 7.79e-4 | 9.79e-4 |  |
| **Model diagnostic tests** | |  |  |  |  |  |
| Test name | n | Observed | Expected | p value |  |  |
| Uniformity | - | 0.0446 | - | 0.0208 |  |  |
| Dispersion | - | 0.9031 | - | 0.5360 |  |  |
| Temporal autocorrelation (survey year) | 76 | 1.6736 | - | 0.2458 |  |  |
| Spatial autocorrelation (reef coordinates) | 23 | -0.0122 | -0.0531 | 0.4305 |  |  |

***Table S2:*** *Pearson's correlation coefficient for anomaly values estimated using Local Outlier Factor set with different neighborhood sizes (k in Supplementary Methods equations).*

|  | Local Outlier Factor neighborhood size | | | | | | |
| --- | --- | --- | --- | --- | --- | --- | --- |
|  | 3 | 4 | 5 | 6 | 7 | 8 | 9 |
| 4 | 0.864 |  |  |  |  |  |  |
| 5 | 0.791 | 0.939 |  |  |  |  |  |
| 6 | 0.747 | 0.877 | 0.943 |  |  |  |  |
| 7 | 0.725 | 0.849 | 0.916 | 0.978 |  |  |  |
| 8 | 0.707 | 0.827 | 0.888 | 0.942 | 0.968 |  |  |
| 9 | 0.671 | 0.772 | 0.827 | 0.865 | 0.895 | 0.938 |  |
| 10 | 0.605 | 0.683 | 0.725 | 0.757 | 0.787 | 0.832 | 0.912 |

**Table S3:** Model summary table for linear regression of maximum reef novelty as a function of the reef-level intercepts and slopes from the predictive anomaly model (Table S1).

| Variable | Estimate | Std Error | *t*-value | *p*-value |
| --- | --- | --- | --- | --- |
| Intercept | 0.135 | 0.007 | 19.977 | < 0.001 |
| Reef-level intercept | 0.049 | 0.010 | 4.793 | < 0.001 |
| Reef-level slope | 0.046 | 0.010 | 4.567 | < 0.001 |
| Intercept:slope | -0.063 | 0.004 | -1.425 | 0.157 |

**Supplementary figures**

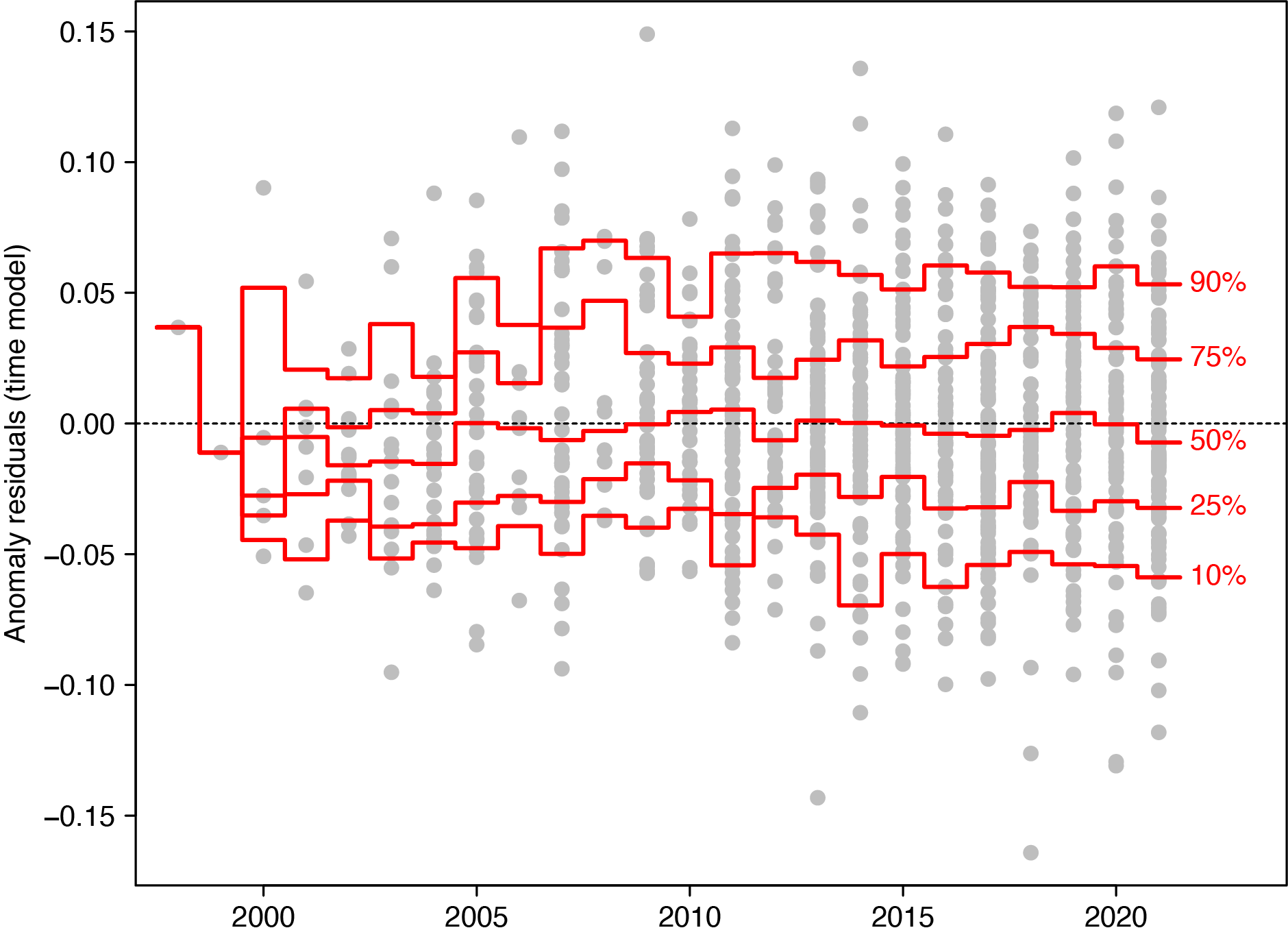

***Figure S1:*** *Quantiles of LOF (novelty) residuals from the full over time model in the main text. Quantiles were calculated for each survey year, with any trend in quantiles suggesting the distribution of anomaly values changed over time. The absence of these trends suggests that anomaly results in the main text link to systematic compositional movement of reefs away from past states, rather than an increase in large-value anomalies.*

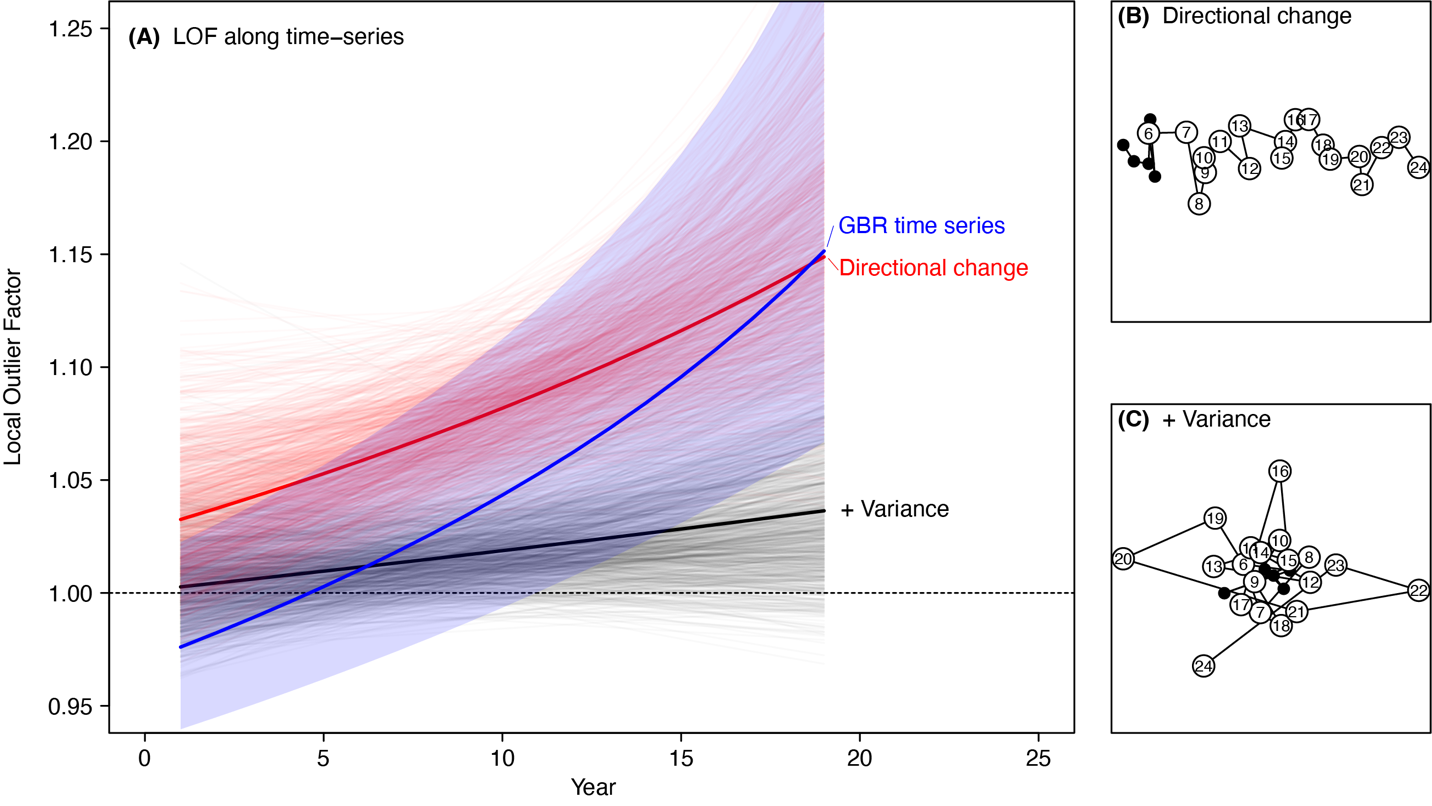

***Figure S2:*** *Local Outlier Factor over time using observed GBR time series (blue), as well as two sets of 999 simulated time series. The directional change simulation (B) replicates the gradual loss of three of ten assemblage taxa over a twenty-year period, creating a strong directional trend in compositional space. The variance simulation (C) increases year-to-year compositional noise in a linear fashion by adding random Gaussian draws to static taxa abundances.*

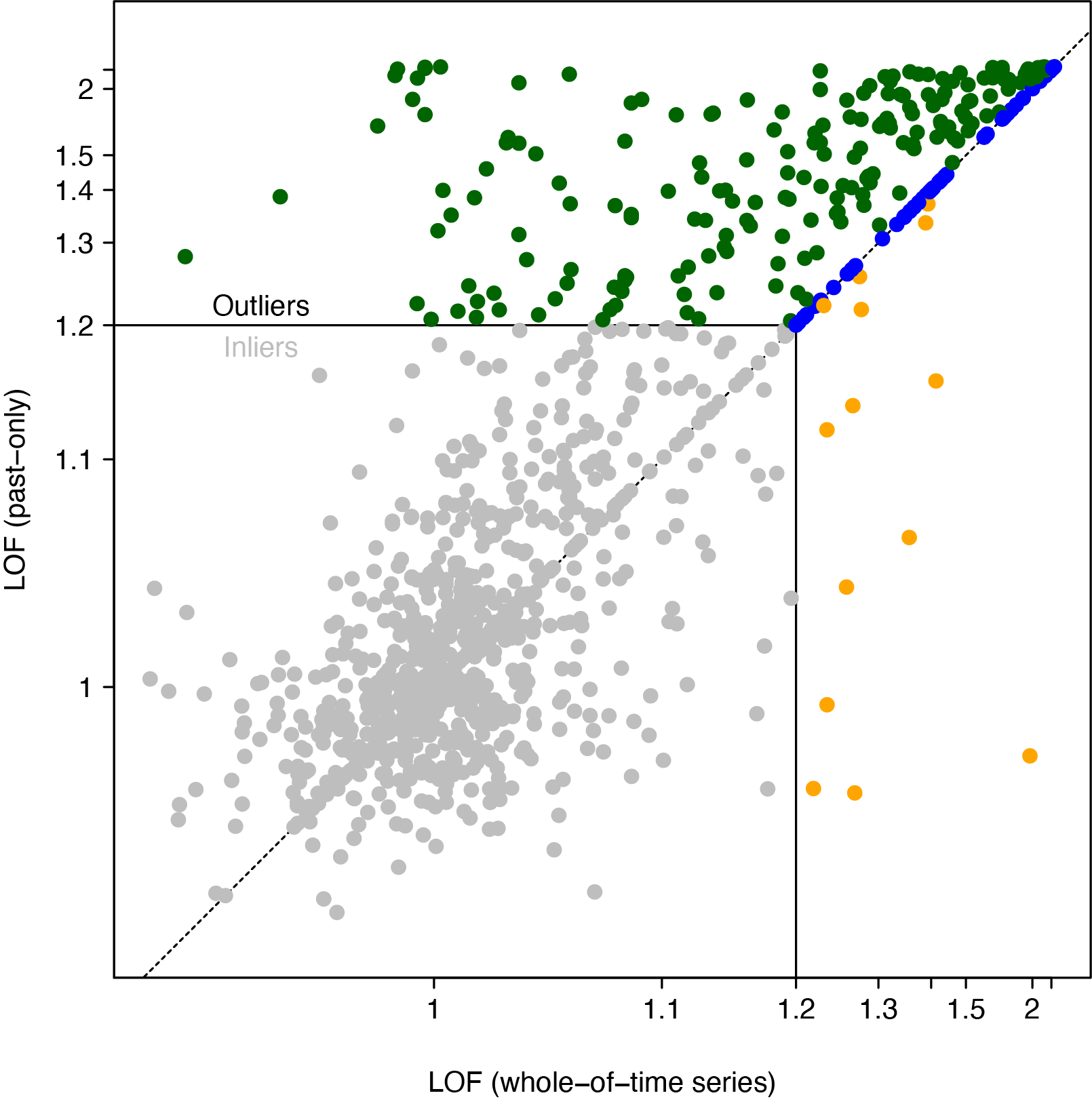

***Figure S3:*** *Comparison of Local Outlier Factors calculated using only past time series surveys (y-axis, "Past-only", as per main text) and calculated using all time series surveys (x-axis, "Whole-of-time series"). Points below 1.2 were considered "inliers", with "outliers" classed into those with a greater (green), same (blue) or lower (yellow) Past-only LOF.*

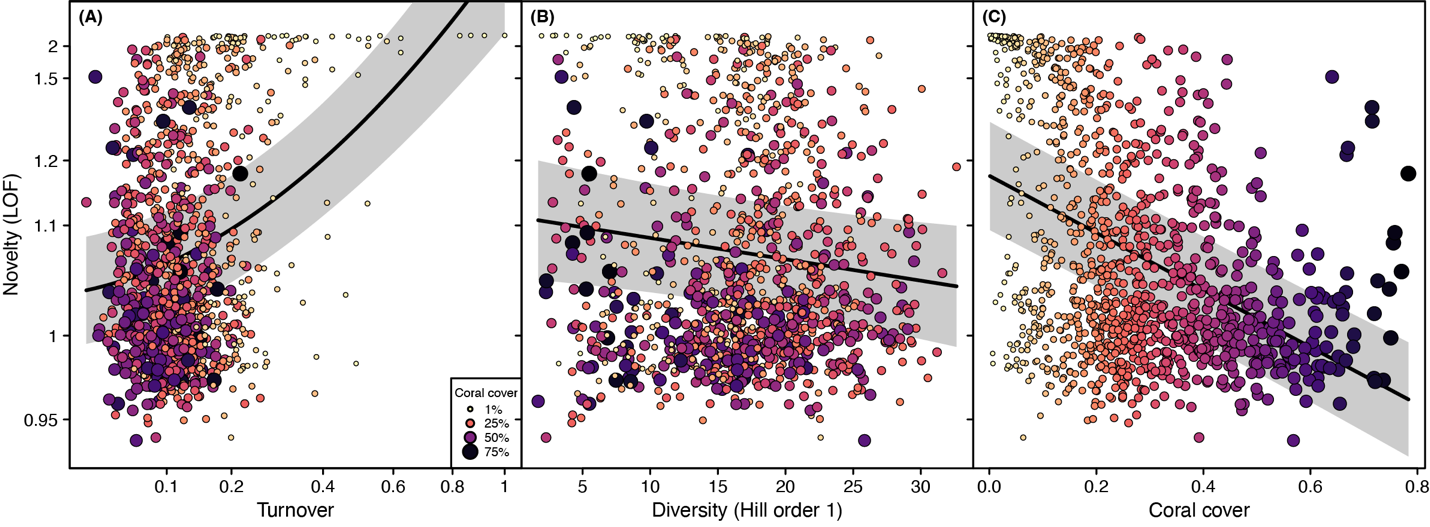
***Figure S4:*** *Correlation of novelty with significant predictors, including (A) Year to year compositional turnover, measured using Bray-Curtis dissimilarity, (B) assemblage diversity as Hill order 1 (exponent of Shannon's diversity) and (C) coral cover. Points sized and coloured by coral cover. Novelty y-axes are Box-Cox transformed.*

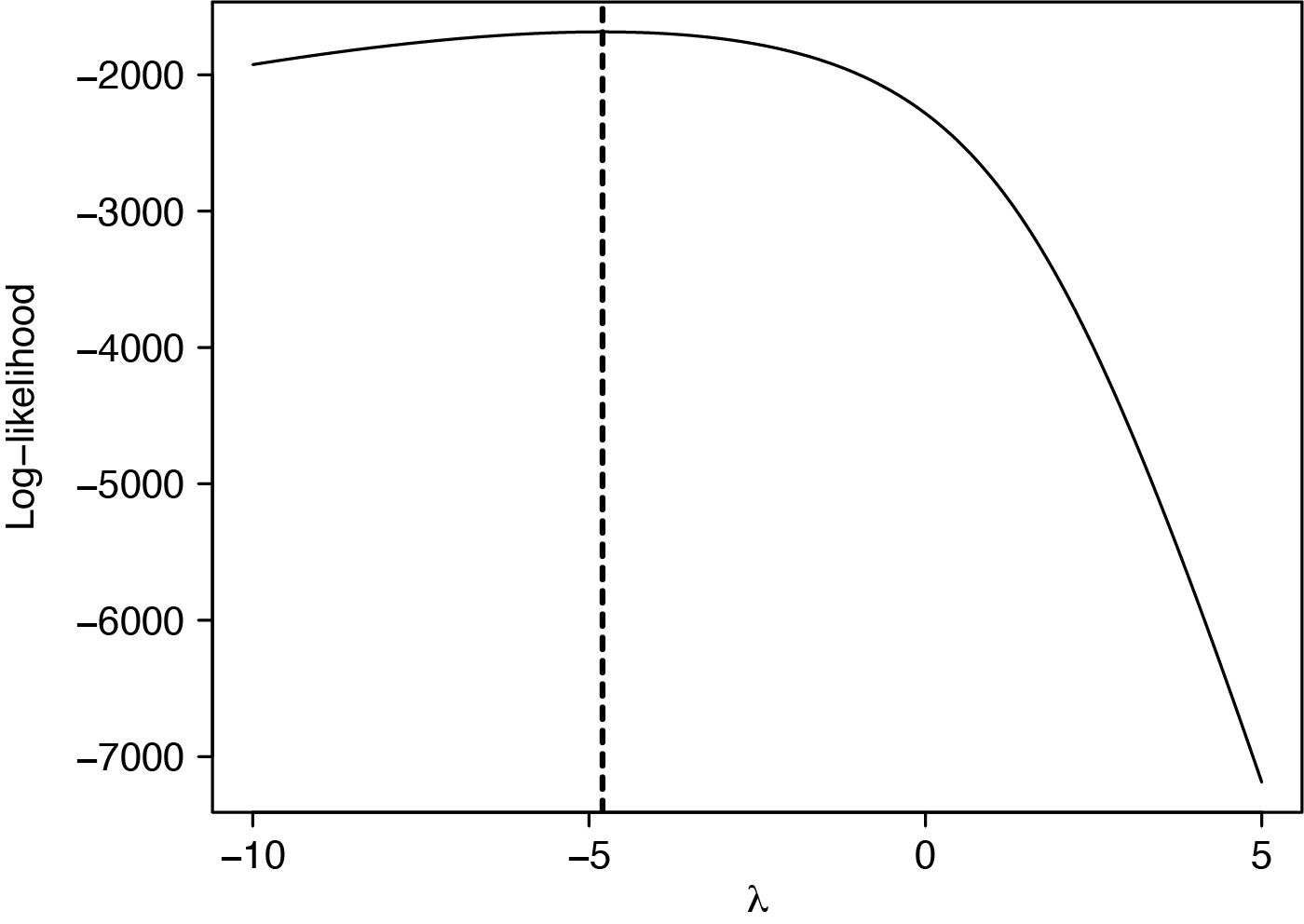

***Figure S5:*** *Log-likelihood test of λ scaling parameter of Box-Cox transformation for Local Outlier Factor novelty estimation.*
